# Large-scale proteomic analysis of human prefrontal cortex identifies proteins associated with cognitive trajectory in advanced age

**DOI:** 10.1101/526988

**Authors:** Aliza P. Wingo, Eric B. Dammer, Michael S. Breen, Benjamin A. Logsdon, Duc M. Duong, Juan C. Troncosco, Madhav Thambisetty, Thomas G. Beach, Geidy E. Serrano, Eric M. Reiman, Richard J. Caselli, James J. Lah, Nicholas T. Seyfried, Allan I. Levey, Thomas S. Wingo

## Abstract

In advanced age, some individuals maintain a stable cognitive trajectory while others experience a rapid decline. Such variation in cognitive trajectory is only partially explained by traditional neurodegenerative pathologies. Hence, to identify new processes underlying variation in cognitive trajectory, we perform an unbiased proteome-wide association study of cognitive trajectory in a discovery (*n*=104) and replication cohort (*n*=39) of initially cognitively unimpaired, longitudinally assessed older-adult brain donors. We find 579 proteins associated with cognitive trajectory after meta-analysis. Notably, we present novel evidence for increased neuronal mitochondrial activities in cognitive stability regardless of the burden of traditional neuropathologies. Furthermore, we provide additional evidence for increased synaptic activities and decreased inflammation and apoptosis in cognitive stability. Importantly, we nominate proteins associated with cognitive trajectory, particularly the 38 proteins that act independently of neuropathologies and are also hub proteins of protein co-expression networks, as promising targets for future mechanistic studies of cognitive trajectory.

---

In advanced age, trajectory of cognitive performance over time can range from stability to rapid decline. Like all complex human traits, cognitive trajectory varies from person to person and is determined by a combination of genetic and environmental factors including premorbid cognitive abilities, educational attainment, life-style choices, and environmental exposures^1 2^. A stable cognitive trajectory in advanced age is desirable and may reflect cognitive resilience, while decline in cognitive performance may be the earliest manifestation of a serious neurodegenerative disease^1 3^. Cognitive decline may ultimately lead to a diagnosis of mild cognitive impairment (MCI) or dementia. Hence, variation in cognitive trajectory can influence onset age for MCI or dementia.

Studying individual cognitive trajectory is strategic for several reasons. First, it captures all the factors affecting cognition, including diverse pathologies as well as biological mechanisms independent of pathologies^4–6^. Second, it likely captures co-occurring disease processes and co-occurring age-related pathologies, which are known to be prevalent in the brains of aged individuals^4 5^. Thus, it provides more comprehensive information than the traditional case/control status based on clinical or pathological diagnosis. Third, cognitive trajectory is estimated from periodically and prospectively assessed cognitive performance. Such trajectory captures the progressive nature of cognitive decline, from the asymptomatic stage to the manifestation of mild cognitive impairment or dementia, and thus is more germane to early intervention and prevention. Fourth, as will be shown, within each category of pathology or clinical diagnosis of cognitive status, different persons have different cognitive trajectories making cognitive trajectory relevant to individuals with or without dementia.

Despite variation in cognitive trajectory, in advanced age the majority of individuals tend to experience cognitive decline. Cognitive decline has been suggested to be attributable to dementia pathologies. However, pathologies such as amyloid plaques, neurofibrillary tangles, microinfarct, macroinfarct, and Lewy bodies together capture only about 41% of the variance in cognitive trajectory, leaving 59% unexplained^5 7^. It is unclear what causes the large majority of the variance in cognitive trajectory. Proteins, as functional gene products, are well-suited for studying biological processes underlying individual variation in cognitive trajectory. Yet, no proteome-wide study of cognitive trajectory has been performed to date.

To identify new biological processes underlying cognitive trajectory, we performed the first unbiased, large-scale proteome-wide association study of cognitive trajectory using a discovery and replication cohort of initially cognitively unimpaired, longitudinally assessed older-adult brain donors (Figure 1). We then performed Gene Ontology enrichment analysis on significantly associated proteins to glean a deeper understanding of the biological processes underlying individual differences in cognitive trajectory. Next, we determined which proteins associated with cognitive trajectory were key drivers of protein co-expression networks because these proteins are likely promising targets for future mechanistic studies of cognitive trajectory. Lastly, we determined the proteins that were associated with cognitive trajectory independently of the traditional Alzheimer’s disease pathologies (Figure 1). Together, our findings established a new framework for understanding molecular mechanisms underlying individual differences in cognitive trajectory.

**Figure 1:**
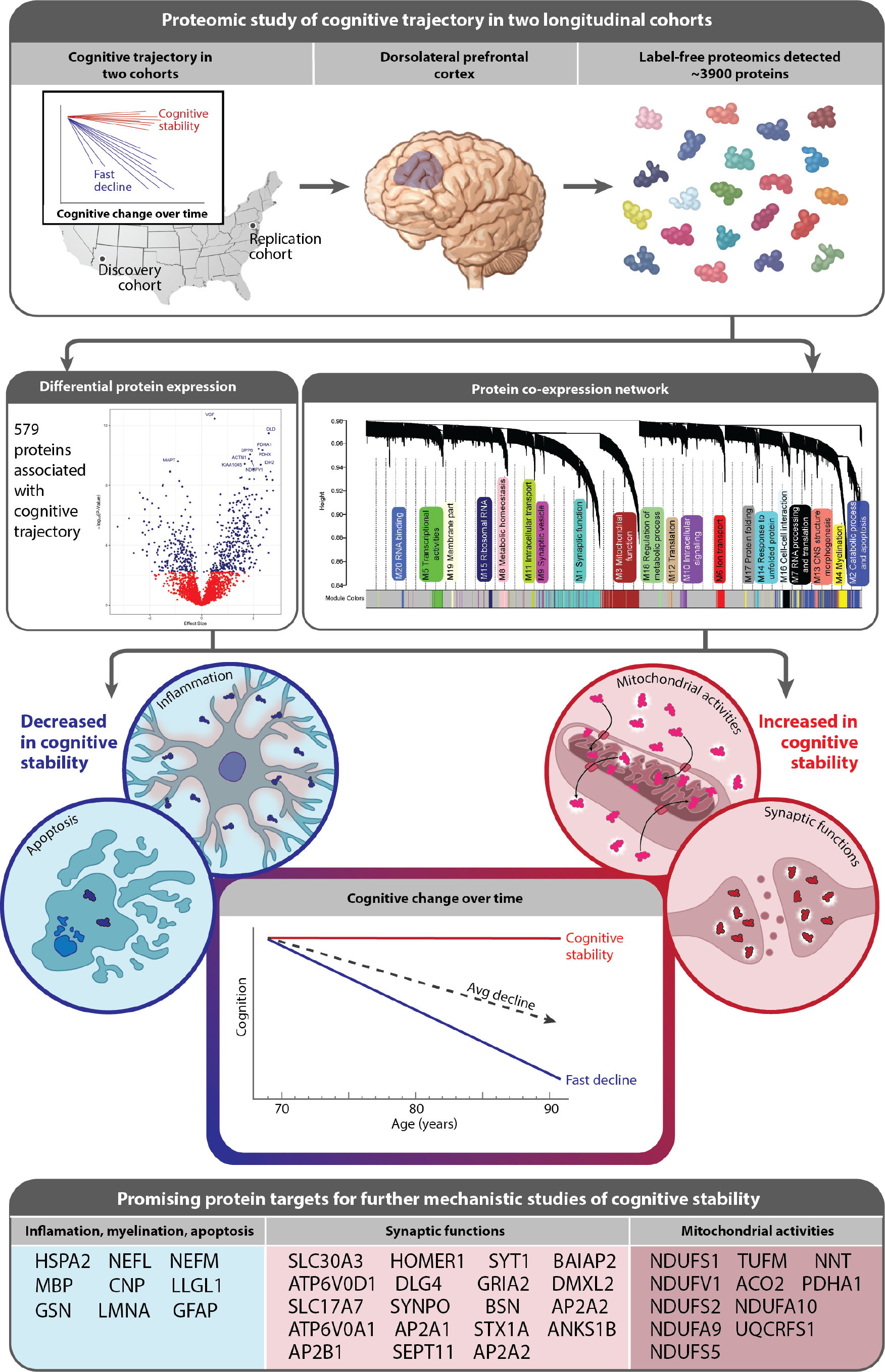
Overview of the study design and results.

## METHODS

### Subjects

#### Banner Sun Health Research Institute participants

This project, the Arizona Study of Aging and Neurodegenerative Disorders, is a longitudinal clinicopathological study of normal aging, Alzheimer’s disease (AD), and Parkinson’s disease (PD)^8^. Most subjects were enrolled as cognitively unimpaired volunteers from the retirement communities of the greater Phoenix, Arizona, USA^8^. Recruitment efforts were also directed at subjects with AD and PD from the community and neurologists’ offices. Subjects received standardized general medical, neurological, and neuropsychological tests annually during life and more than 90% received full pathological examinations after death^8^. For this study, we only included subjects with a clinical diagnosis of AD or normal control rendered approximate to death. Please refer to Table 1 for subject characteristics. Additional inclusion criteria for this study were having a baseline score from the Mini Mental State Examination (MMSE) of 27 or above (i.e. non-demented at baseline)^9 10^ and at least one additional MMSE score in subsequent follow-up visits.

**Table 1:** Demographic and clinical characteristics of the discovery and replication cohorts

| **Characteristic** | **Banner**<br>**(N=104)** | **BLSA**<br>**(N=39)** |
| --- | --- | --- |
| **Age at enrollment**<br>Mean ± SD<br>Median; Range | 80.7 ± 6.6<br>80.6; [61.5 – 96.4] | 74.8 ± 6.9<br>75.3; [58.5 - 89.3] |
| **Age at death**<br>Mean ± SD<br>Median; Range | 87.0 ± 6.7<br>87; [74 – 103] | 87.2 ± 7.7<br>91; [71 – 99] |
| **Sex** (% female, N) | 48.1% (50) | 38.5% (15) |
| **Education** (years)<br>Mean ± SD<br>Median; Range | 14.6 ± 2.6<br>16; [8.0 – 20.0] | 16.6 ± 3.0<br>18; [8.0 – 20.0] |
| **Clinical diagnosis**<br>AD (% , N)<br>Control (% , N) | 26.9% (28)<br>73.1% (76) | 41.0% (16)<br>59.0% (23) |
| **Follow-up duration** (years)<br>Mean ± SD<br>Median; Range | 5.1 ± 3.2<br>4.4; [0.5 – 14.0] | 9.3 ± 3.9<br>8.5; [3.9 – 20.4] |
| **PMI** (h)<br>Mean ± SD<br>Median; Range | 2.9 ± 0.9<br>2.7; [1.2 – 5.5] | 13.8 ± 5.5<br>14.5; [2.0 – 24.0] |
| **Braak** (% , N)<br>I<br>II<br>III<br>IV<br>V<br>VI | 4.8% (5)<br>10.6% (11)<br>25.0% (26)<br>39.4% (41)<br>14.4% (15)<br>5.8% (6) | 0% (0)<br>15.4% (6)<br>15.4% (6)<br>33.3% (13)<br>10.3% (4)<br>25.6% (10) |
| **CERAD** | Criteria not met: 12.5% (13)<br>Definite AD: 25.0% (26)<br>No AD: 28.8% (30)<br>Possible AD: 31.7% (33)<br>Probable AD: 1.9% (2) | 0 (none): 20.5% (8)<br>A (sparse): 7.7% (3)<br>B (moderate): 30.8% (12)<br>C (frequent): 41.0% (16) |
| **Slope of cognitive trajectory (MMSE/year)**<br>*All participants*<br>Mean ± SD<br>Median; Range<br>*By clinical diagnosis (median; range)*<br>Controls<br>AD cases<br>*By Braak stage (median; range)*<br>I and II<br>III and IV<br>V and VI | -0.5 ± 0.8<br>-0.3; [-4.9 to 0.2]<br>-0.1; [-1.6 to 0.2]<br>-1.1; [-4.9 to -0.1]<br>-0.1; [-0.9 to 0.2]<br>-0.2; [-2.1 to 0.1]<br>-1.1; [-4.9 to -0.1] | -0.4 ± 0.6<br>-0.1; [-2.4 to 0.1]<br>-0.1; [-0.7 to 0.1]<br>-0.4; [-2.4 to 0.0]<br>-0.2; [-0.7 to 0.0]<br>0.0; [-2.4 to 0.1]<br>-0.4; [-1.8 to 0.0] |

AD: Alzheimer's disease; PMI: post-mortem interval
Braak: neurofibrillary tangle stages. Cerad: severity of amyloid plaques.

#### Baltimore Longitudinal Study of Aging (BLSA) participants

BLSA is a prospective cohort study of aging in community dwelling individuals^11 12^. It continuously recruits healthy volunteers aged 20 or older and follow them for life regardless of changes in health or functional status. Participants were examined at the National Institute of Aging Clinical Research Unit in Baltimore at one to four-year intervals, with more frequent follow-up visits for older participants^11 12^. Due to a smaller sample size and the fact that these subjects were recruited as cognitively unimpaired individuals, all BLSA individuals with proteomic data were included regardless of MMSE score at baseline.

### Personalized cognitive trajectory

Person-specific cognitive trajectory was estimated using a linear mixed model for each subject. For each cohort, we model the annual MMSE score as a longitudinal outcome, follow-up year as the independent variable and sex, age at enrollment, and education as the covariates, with a random intercept and random slope per subject using the lme4 R package (version 1.1-19). The derived cognitive trajectory for each individual in Banner and BLSA cohorts is listed in supplementary table 1.

### Pathological assessment of Banner and BLSA cohorts

Measures of AD pathology, i.e., beta-amyloid plaques and neurofibrillary tangles, were used as covariates to adjust for the degree of AD pathology in each sample. Beach et al. described in detail how these measures were derived for the Banner cohort^8^. Briefly, plaque total is the summary density score of all types of amyloid plaques in the frontal, temporal, and parietal cortices, as well as hippocampus CA1 region and entorhinal region. Additionally, neuritic plaque density scoring was done according to the Consortium to Establish a Registry for Alzheimer’s Disease (CERAD)^10^. Tangle total is the sum of neurofibrillary tangle density in the frontal cortex, temporal cortex, parietal cortex, hippocampus, and entorhinal region. Tangle scoring was done according to the CERAD templates^10^. In the BLSA cohort, O’Brien et al. described how estimates of beta-amyloid positive plaques and neurofibrillary tangles were derived^13^. Briefly, silver staining was performed to assess severity of neuritic plaques using CERAD (score ranges from 0 to 3) and of neurofibrillary tangles using Braak (score ranges from 1 to 6), with higher scores indicating higher level of severity.

### Proteomic Quantification

#### Brain Tissue Homogenization and Proteolytic Digestion

Whole-brain proteomes were derived from cortically dissected dorsolateral prefrontal cortex for the discovery and replication cohorts, separately. All tissues were homogenized in 8M Urea buffer and digested with trypsin as previously described^14^. In the Banner cohort, samples were randomized with respect to sex, age at death, diagnosis, race, and *APOE* genotype. In BLSA cohort, samples were randomized with respect to sex, age at death, diagnosis, and *APOE* genotype.

#### Liquid Chromatography Coupled to Tandem Mass Spectrometry (LC-MS/MS)

Proteomic quantification for both cohorts used the approach described in Seyfried et al.^14^, and a complete description of the methods and all raw proteomic data from the Banner and BLSA cohorts are given in these references^15 16^. Briefly, brain-derived tryptic peptides were separated using a NanoAcquity UHPLC (Waters, Milford, FA) and monitored on a Q-Exactive Plus mass spectrometer (ThermoFisher Scientific, San Jose, CA). Raw data were analyzed using MaxQuant v1.5.2.8 with Thermo Foundation 2.0 for RAW file reading capability^17^. Co-fragmented peptide search was enabled to deconvolute multiplex spectra. The false discovery rate (FDR) for peptide spectral matches, proteins, and site decoy fractions were set to 1%. Protein quantification was estimated by label free quantification (LFQ) algorithm by MaxQuant and only considered razor plus unique peptides for each protein isoform. In essence, this approach estimated the protein abundance for a given protein isoform by using peptides that are unique to the specific isoform and peptides that map to multiple isoforms of the same gene. In our prior work on the BLSA dataset, we used data from the dorsolateral prefrontal cortex (dPFC) and precuneus regions together; however, in this work, we only focused on proteins from the dPFC. After label-free quantification, 3710 proteins in Banner and 3933 proteins in BLSA cohorts were detected, and 2752 protein isoforms were detected in both Banner and BLSA cohorts.

#### Proteomic Quality Control

Only proteins quantified in at least 90% of the samples were included in the analysis. We performed log2 transformation of the proteomic profiles. Subsequently, linear mixed models from the R package variancePartition^18^ were used to characterize the percent of protein variance explained by biological and technical variables. This approach was used to quantify the main sources of variation in each proteomic dataset separately and identified the variance attributable to age, sex, neuropathology, post-mortem interval (PMI), and batch. For instance, in the Banner cohort, batch and sex explained substantial proportions while age at death and PMI explained some proportions of variance of the proteomic profile (Supplementary Figure 1). Therefore, within each cohort, effect of batch was removed using Combat^19^, and effects of sex, age at death, and PMI were removed using bootstrap regression. We removed these technical (batch and PMI) and biological effects (sex and age at death) because they may confound our association analyses between cognitive trajectory and protein levels. Supplementary Figure 2 shows that after Combat and bootstrap regression, batch, sex, age at death, and PMI explained only minimal amounts of the variance of the proteomic profile. The same approach was used to successfully remove the effects of these same covariates from the BLSA proteome^14^.

### Bioinformatics

#### Proteome-wide association study of cognitive trajectory

Proteome-wide association analysis was performed in each cohort separately, and meta-analysis was used to combine individual results. In each cohort, linear regression was performed with cognitive trajectory as the outcome and normalized protein abundance as the predictor. In our planned secondary analysis, we performed the same association analysis as above but adding beta-amyloid plaques and neurofibrillary tangles as covariates. Meta-analysis was performed with METAL^20^, a popular meta-analysis tool^21 22^, using effect size estimates and standard errors from the proteome-wide association studies of cognitive trajectory in Banner and BLSA, respectively. We included proteins that are detected in both the Banner and BLSA cohorts in the meta-analysis. For all analyses, we used Benjamini-Hochberg (BH) method to control the false discovery rate (FDR)^23^, and declared significantly associated proteins as those with BH FDR p<0.05.

#### Protein co-expression network analysis

Network analysis was performed in Banner and BLSA datasets separately to identify modules of co-expressed proteins. For the Banner network, missing proteins were imputed using the k-nearest neighbor imputation function in R. Then, batch effects were removed using Combat^19^, and age at death, sex, and PMI were regressed from the proteomic profiles using Bootstrap regression. Weighted gene co-expression network analysis (WGCNA)^24^ was used on normalized protein abundance to define protein co-expression networks. For BLSA, we used the BLSA networks from Seyfried et al.^14^, which were previously built using proteins measured from the precuneus and prefrontal cortex in the same individual. We defined hub proteins, i.e. highly connected proteins, for each of the modules as those with intramodular kME in the top 90^th^ percentile among the proteins in the corresponding module^24^. Gene ontology (GO) enrichment analysis was performed on each protein co-expression module using GO Elite and Fisher exact test^25^ to glean a deeper biological understanding of these modules.

#### Enrichment analysis of proteins associated with cognitive trajectory

GO enrichment analysis was performed on the proteins associated with cognitive trajectory at FDR <0.05 from the meta-analysis. Proteins were categorized into two lists for GO enrichment analysis: i) higher abundance in cognitive stability, and ii) lower abundance in cognitive stability. Gene sets were downloaded from MSigDB (version 2017), including GO biological process, cellular component, and molecular function. We also downloaded the cell type-specific protein expression in the brain generated by Sharma et al^26^, as well as cell type-specific gene expression data generated by Darmanis et al^27^ and Zeisel et al^28^ to assess for enrichment of the significantly associated proteins/genes in particular brain cell types. For enrichment analysis, we used a modified version of the GeneOverlap function (from package of the same name) in R and Fisher’s exact test so that all pairwise tests were adjusted for multiple testing using BH method to control the FDR^23^. The Fisher’s test function also provides an estimated odds-ratio in comparison to a proteomic background set to 20,000.

##### Data availability

All proteomic and phenotypic data that support the findings of this study are contained in Supplementary Table 1 and in https://www.synapse.org/#!Synapse:syn7170616 and https://www.synapse.org/#!Synapse:syn3606086.

## RESULTS

### Subjects

The Banner discovery cohort consisted of 104 participants who were non-demented at baseline and were followed for up to 14 years (Table1). Among these, 48% were women, 27% had the final clinical diagnosis of AD, 73% had the final diagnosis of normal control, and median age at death was 87 (Table 1). The BLSA replication cohort had 39 participants who were followed for up to 20 years (Table 1). Of these, 39% were women, 41% had a final clinical diagnosis of AD and 59% had a final clinical diagnosis of normal control, and median age at death was 91 (Table 1).

### Personalized cognitive trajectories

Individual cognitive trajectory was estimated for each participant in both cohorts (Figure 2; Table 1). Cognitive trajectories by last clinical diagnosis before death and by Braak score were depicted in Figure 2. Trajectories with slopes at or near zero reflect cognitive stability, while trajectories with large negative slopes indicate faster cognitive decline. We depicted cognitive trajectory by Braak score for demonstration here because tangles have been suggested to account for more cognitive decline than plaques^29^. The median rate of cognitive change was −0.3 unit of MMSE score per year for Banner participants and −0.1 unit of MMSE per year for BLSA participants (Table 1). Analyzed jointly, the correlation between cognitive trajectory and clinical diagnosis of cognitive status was −0.49 (95% confidence interval (CI) of −0.60 to −0.35, p-value of 4.3×10^−10^) and between cognitive trajectory and Braak score was −0.39 (95% CI of −0.51 to −0.24, p-value of 1.5×10^−6^). These data reveal that i) cognitive trajectory varied within a given clinical diagnosis or Braak score; ii) faster cognitive decline was not exclusively seen in AD cases or in brains with greater Braak scores; and, iii) cognitive trajectory provides complementary information to the clinical diagnosis and neuropathology findings.

**Figure 2:**
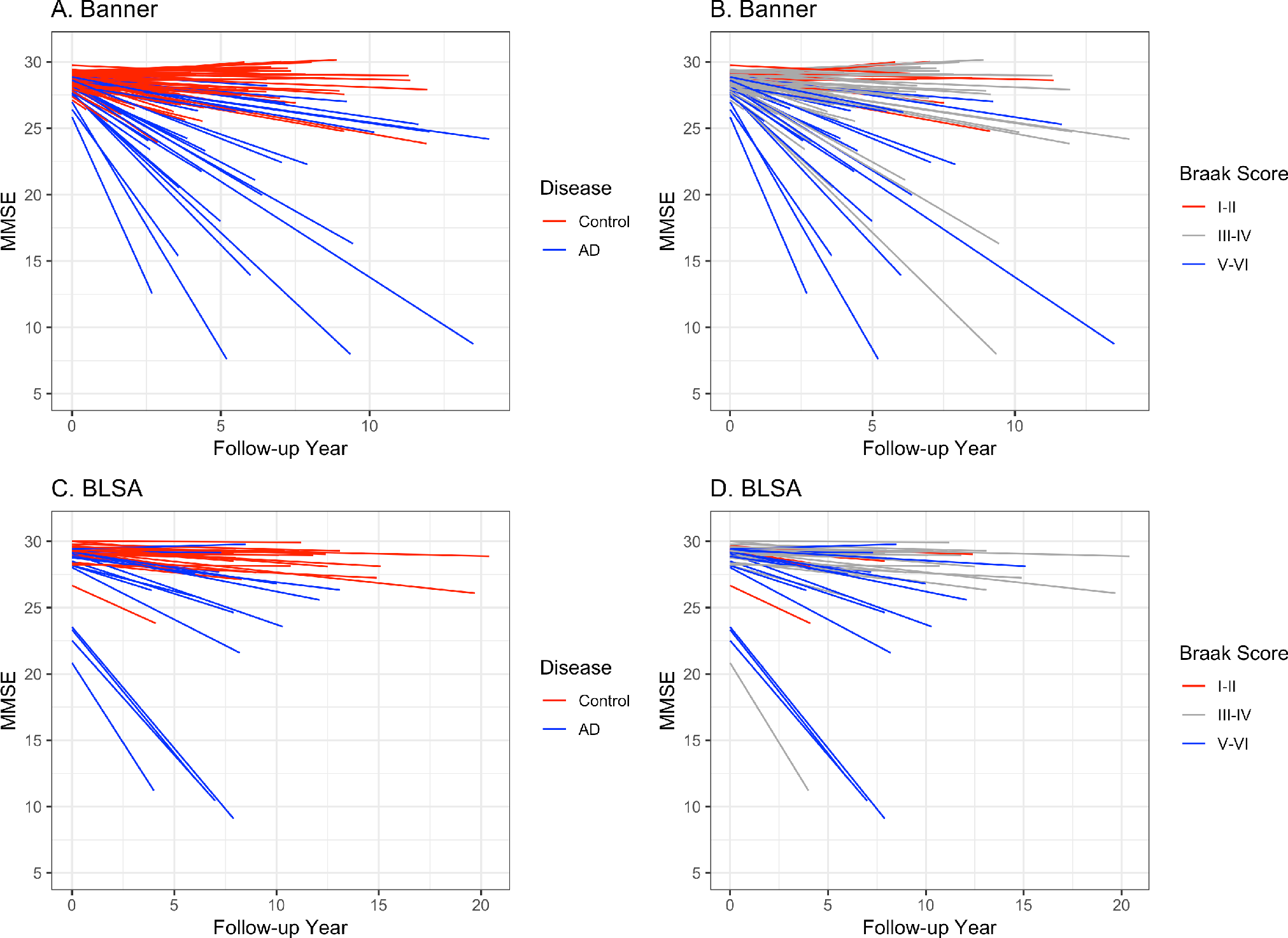
Person-specific cognitive trajectory by clinical diagnosis and Braak score in Banner and BLSA cohorts. Individual cognitive trajectory in the Banner cohort by **a)** last clinical diagnosis (AD or control) prior to death; **b)** Braak score. Individual cognitive trajectory in the BLSA cohort by **c)** clinical diagnosis; **d)** Braak score. A positive slope or small negative slope reflects cognitive stability. A larger negative slope reflects faster cognitive decline.

### Proteome-wide association study of cognitive trajectory

After label-free quantification, 3710 proteins in Banner cohort and 3933 proteins in BLSA cohort were detected with 10% or fewer missing values. Among the detected proteins, 2752 protein isoforms were detected in both Banner and BLSA cohorts. In the Banner discovery cohort, 354 proteins were associated with cognitive trajectory at FDR <0.05 after adjusting for sex, age at enrollment, education, age at death, PMI, and batch (N=104, Supplementary Table 1a; Supplementary Figure 2A). The top five proteins were VGF, SEPT5, DBI, MAPT, and KIAA1045 (aka PHF24). In the BLSA replication cohort, 127 proteins were associated with cognitive trajectory at FDR <0.05 after adjusting for sex, age at enrollment, education, age at death, and PMI (N=39; Supplementary Table 2, Supplementary Figure 2B). The top 5 proteins were DLD, ABHD10, VDAC1, NDUFV1, and PDHB. Likely reasons for difference in the top differentially expressed proteins in the two cohorts are the differences in demographic composition, follow-up duration, range of cognitive trajectory, and false-discovery. Given these differences, a meta-analysis of these data was performed to enhance true positive findings, as the meta-analysis is designed to integrate the results of several independent studies and provide a more precise estimate of the association between predictor and outcome than any individual study contributing to the pooled analysis^30^.

The meta-analysis showed 579 proteins associated with cognitive trajectory in the same directions in both the discovery and replication cohorts and at FDR <0.05 among the 2752 protein isoforms commonly detected in both cohorts (N=143 individuals; Figure 3A, supplementary table 3). Regarding the direction of association, 350 proteins had increased abundance while 229 had decreased abundance in cognitive stability. For instance, VGF protein had increased abundance and MAPT protein had decreased abundance in cognitive stability (Figures 3B,C). VGF is a neuropeptide that regulates synaptic function^31^, synaptic plasticity^32^, and hippocampal memory consolidation^33^. MAPT (microtubule associated protein tau) is involved in axonal transport, synaptic plasticity, and synaptic function^34^, and aggregation of hyperphosphorylated tau proteins can lead to formation of neurofibrillary tangles and Alzheimer’s dementia^35^. Going forward, we will refer to proteins with increased abundance in cognitive stability as higher-abundance and proteins with decreased abundance in cognitive stability as lower-abundance proteins.

**Figure 3:**
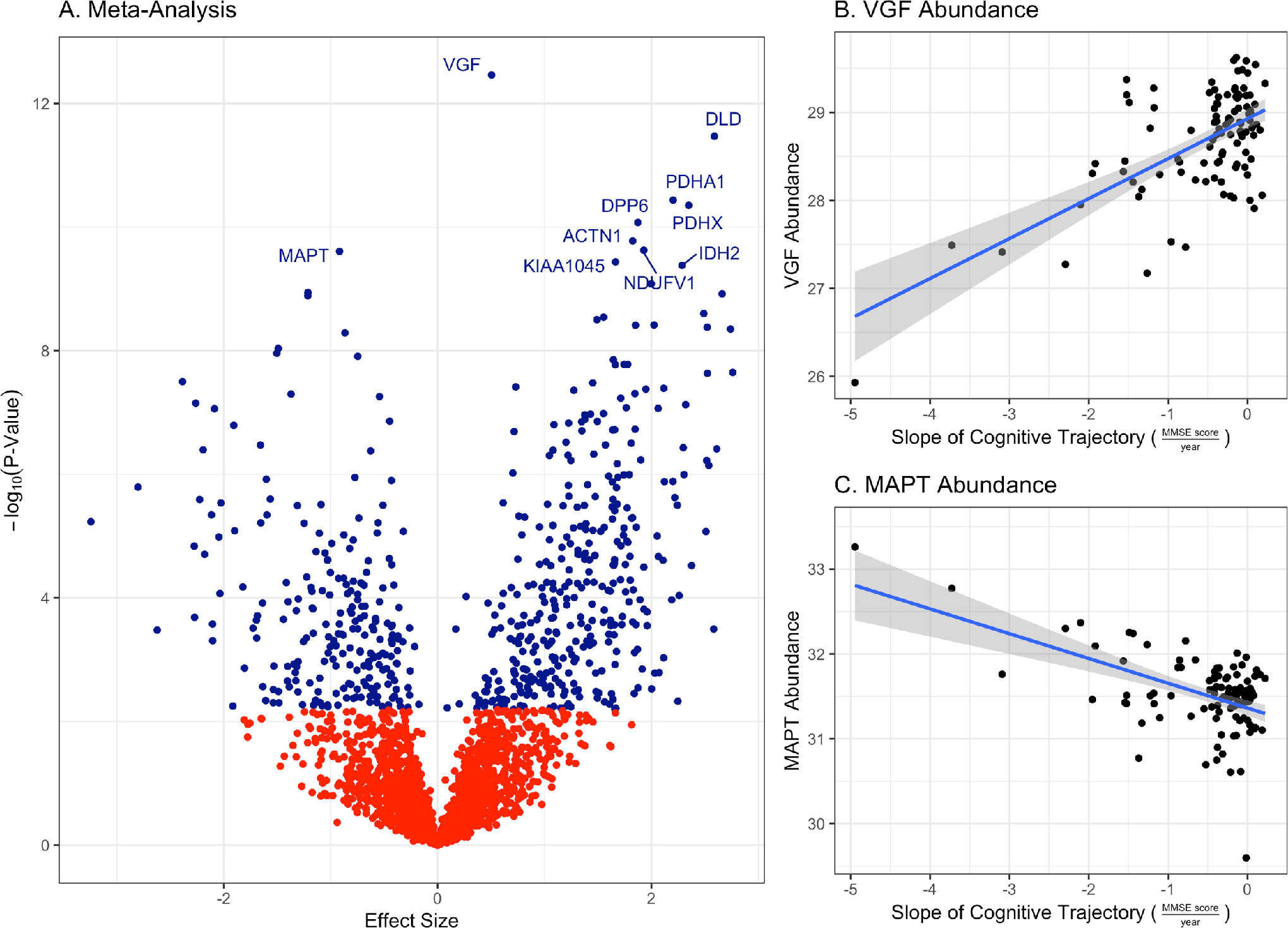
**a)** Volcano plot for meta-analysis of proteome-wide association studies of cognitive trajectory in the discovery and replication cohorts. A total of 579 proteins were associated with cognitive trajectory at FDR <0.05. Among these, 350 proteins had increased abundance and 229 had decreased abundance in cognitive stability; **b)** Higher VGF protein level was associated with cognitive stability. Of note, for cognitive trajectory, a small negative slope reflects slow decline and a large negative slope reflects fast decline. Cognitive stability is reflected by a positive slope or very small negative slope of the cognitive trajectory. The grey area represents 95% confidence interval for the best-fit line between cognitive trajectory and VGF abundance; **c)** Lower MAPT protein level was associated with cognitive stability.

Of the 579 cognitive trajectory-associated proteins, we hypothesized that some would interact potentially as members of distinct biological processes or organelles. Thus, we asked whether there was evidence for protein-protein interaction (PPI) using experimentally validated data available on BioGRID v3.5.165^36^. The 569 unique proteins among these 579 proteins (10 had isoforms) were categorized into presynaptic (N=159), postsynaptic (N=150), both (present in both presynaptic and postsynaptic density; N=20), or other (N=240) using Boyken et. al. presynaptic list^37^ and Bayes et. al. postsynaptic density list^38^ (Supplementary Table 3b). Additionally, we labeled any of these proteins as mitochondrial if they are located in mitochondria based on Uniprot subcellular location^39^. Thus, some proteins may have more than one designation. For instance, of the 157 proteins designated as mitochondrial, 112 were presynaptic, 10 postsynaptic, and 35 were other (Supplementary Table 3b). For the 569 unique proteins, 484 had at least one PPI, and there were a total of 2587 PPIs among these 484 proteins (Supplementary Table 3c; Supplementary Figure 4). Focusing on PPIs emanating from presynaptic proteins, we found 374 proteins with interactions (nodes) and 1745 PPIs (edges) (Supplementary Figure 5A). Focusing on PPIs originating from postsynaptic proteins, we found 353 proteins with interactions (nodes) and 864 PPIs (edges) (Supplementary Figure 5B). Thus there is ample evidence that the proteins associated with cognitive trajectory interact with each other and reveal many PPIs within and between presynaptic and postsynaptic proteins associated with cognitive trajectory. Interestingly, mitochondrial proteins were featured prominently among cognitive trajectory-associated proteins, and more findings about mitochondrial proteins are followed.

### Enrichment analysis of proteins associated with cognitive trajectory

To understand the biological processes related to the 579 proteins associated with cognitive trajectory, we performed GO enrichment analysis for the 229 proteins with lower abundance in cognitive stability and 350 proteins with higher abundance in cognitive stability, separately. The lower-abundance proteins were enriched for inflammatory response, apoptosis, endothelial function, and RNA processing at adjusted p-value <0.05 (Table 2, Supplementary Tables 4a,b,c). The higher-abundance proteins were enriched for mitochondrial function and synaptic transmission at adjusted p-value <0.05 (Table 2, Supplementary Tables 4d,e,f). Next, we mapped these 579 cognitive trajectory-associated proteins to cell types using brain cell type signatures from Sharma et al.^26^, Zeisel et al.^28^, and Damarnis et al.^27^. The lower-abundance proteins in cognitive stability were significantly enriched for signatures of oligodendrocyte, astrocyte, and microglia after multiple testing correction (Table 2, Supplementary Table 5). The higher-abundance proteins were strongly enriched for neurons and mildly enriched for astrocytes at adjusted p-value <0.05 (Table 2, Supplementary Table 5). Together, our findings suggest that the 350 proteins with increased abundance in cognitive stability participate in mitochondrial activities and synaptic transmission in neurons.

**Table 2:** GO enrichment analysis and cell-type enrichment analysis for the 579 cognitive trajectory­ associated proteins. Enrichment analysis was performed separately for the 229 lower-abundance proteins in cognitive stability and 350 higher-abundance proteins in cognitive stability, respectively. Cell type specific signatures were downloaded from Sharma et al., Zeisel et al., and Damarnis et al. Detailed results for these analyses are in Supplementary Tables 4a-f, and 5.

| Low-abundance proteins in Cognitive Stability |  |
| --- | --- |
| GO Biological Process | Inflammatory response; Apoptosis; Endothelial function; RNA processing |
| GO Cellular Component | Inflammatory response; Cellular adhesion |
| GO Molecular Function | RNA binding; Cell-cell adhesion |
| Brain cell type | Oligodendrocyte; Astrocyte; Microglia |
| High-abundance proteins in Cognitive Stability |  |
| GO Biological Process | Mitochondrial function; Synaptic transmission |
| GO Cellular Component | Mitochondrion; Synaptic vesicle; Neuron |
| GO Molecular Function | Cellular respiration; Syntaxin binding |
| Brain cell type | Neuron; Astrocyte |

### Proteome-wide association study of cognitive trajectory adjusting for brain pathology

To determine whether the proteins associated with cognitive trajectory act through or independently of traditional AD pathologies, we performed another proteome-wide association study of cognitive trajectory adjusting for beta-amyloid and tangle pathology in the discovery and replication cohort separately, followed by a meta-analysis. At FDR <0.05, 57 proteins in Banner and 12 proteins in BLSA were associated with cognitive trajectory after adjusting for sex, age at enrollment, education, age at death, PMI, plaques, and tangles (Supplementary Tables 6 and 7). The meta-analysis revealed 232 proteins associated with cognitive trajectory at FDR <0.05 in the same directions in both datasets (Supplementary Table 8).

To identify biological processes associated with these 232 proteins, GO enrichment analysis was performed on the higher-abundance and lower-abundance proteins in cognitive stability separately. Lower-abundance proteins were enriched for the processes of glycolysis, assembly of neurofilament bundle, cell-cell adhesion, and RNA binding, and for oligodendrocyte, astrocyte, and microglia (Table 3; Supplementary Tables 9a-c and 10). Higher-abundance proteins in cognitive stability were enriched for processes of mitochondrial activities and synaptic transmission, and for neurons (Table 3; Supplementary Tables 9d-f and 10).

**Table 3:** GO enrichment analysis and cell type enrichment analysis for the 232 cognitive trajectory­ associated proteins after adjusting for plaques and tangles and other covariates. Enrichment analysis was performed separately for the 81 low-abundance proteins in cognitive stability and 151 high­ abundance proteins in cognitive stability, respectively. Cell type specific signatures were downloaded from Sharma et al., Zeisel et al., and Darmanis et al. Detailed results for these analyses are in Supplementary Tables 9a-f, and 10.

| Low-abundance Proteins in Cognitive Stability after adjusting for plaques and tangles |  |
| --- | --- |
| GO Biological Process | Glycolysis; Neurofilament bundle assembly |
| GO Cellular Component | Focal adhesion; Axon; Neurofibrillary tangle; Microtubule |
| GO Molecular Function | RNA binding; Cell-cell adhesion |
| Brain cell type | Oligodendrocyte; Astrocyte; Microglia |
| High-abundance Proteins in Cognitive Stability after adjusting for plaques and tangles |  |
| GO Biological Process | Mitochondrial activities; Synaptic transmission |
| GO Cellular Component | Mitochondrion; Synaptic vesicle |
| GO Molecular Function | Syntaxin binding |
| Brain cell type | Neuron |

Notably, the proteome-wide association study reveals that the proteins with increased abundance in cognitive stability were involved in mitochondrial activities and synaptic transmission in neurons independently of effects of amyloid plaques and neurofibrillary tangles. In addition, the proteins with decreased abundance in cognitive resilience were involved in inflammatory response, apoptosis, and endothelial function in glial cells, likely acting via plaques and tangles. In contrast to the proteins with higher-abundance in cognitive stability, adjusting for traditional neuropathology influenced those with lower abundance in cognitive stability to a much greater degree and changed biological processes represented.

### Protein co-expression network construction

Many biological functions require a coordinated effort of a large number of genes and proteins, and systems biology approaches seek to identify gene products that act in concert with one another in co-expression networks^40^. We identified 20 modules of strongly co-expressed proteins in the Banner cohort using WGCNA and then gleaned the biological processes enriched in these modules with GO enrichment analysis (Figure 4; Supplementary Tables 11-13). The five largest modules were enriched for synaptic function (M1, 359 proteins), catabolic process and apoptosis (M2, 296 proteins), mitochondrial function (M3, 276 proteins), myelination (M4, 141 proteins), and transcriptional activities (M5, 83 proteins) at adjusted p <0.05 (Supplementary Tables 12-13). Of note, our group previously performed a detailed co-expression network analysis for the BLSA cohort and identified 16 networks of co-expressed proteins^14^. We used these BLSA protein networks in our subsequent analyses.

We performed a pair-wise overlap analysis between pairs of Banner and BLSA protein co-expression modules using the Fisher Exact Test. We found nine pairs of significant overlap after multiple testing correction: Banner M1 (synaptic function) with BLSA M1 (synaptic transmission, ∩=187 proteins, p=2.53E−23, adjusted p=7.40E−06) and BLSA M4 (synaptic membrane/dendrite, ∩=59, p=3.96E−06, adjusted p=0.0015); Banner M2 (catabolic process and apoptosis) with BLSA M2 (myelination, ∩=66, p=4.27E−06, adjusted p=0.0017) and BLSA M6 (inflammatory response, ∩=56, p=2.28E−08, adjusted p=1.54E−05); Banner M3 (mitochondrial function) with BLSA M3 (mitochondrial function, ∩=167, p=5.51E−37, adjusted p=7.40E−06); Banner M4 (myelination) with BLSA M2 (myelination, ∩=100, p=5.96E-17, adjusted p=7.40E−06), among others (Supplementary Table 14).

**Figure 4:**
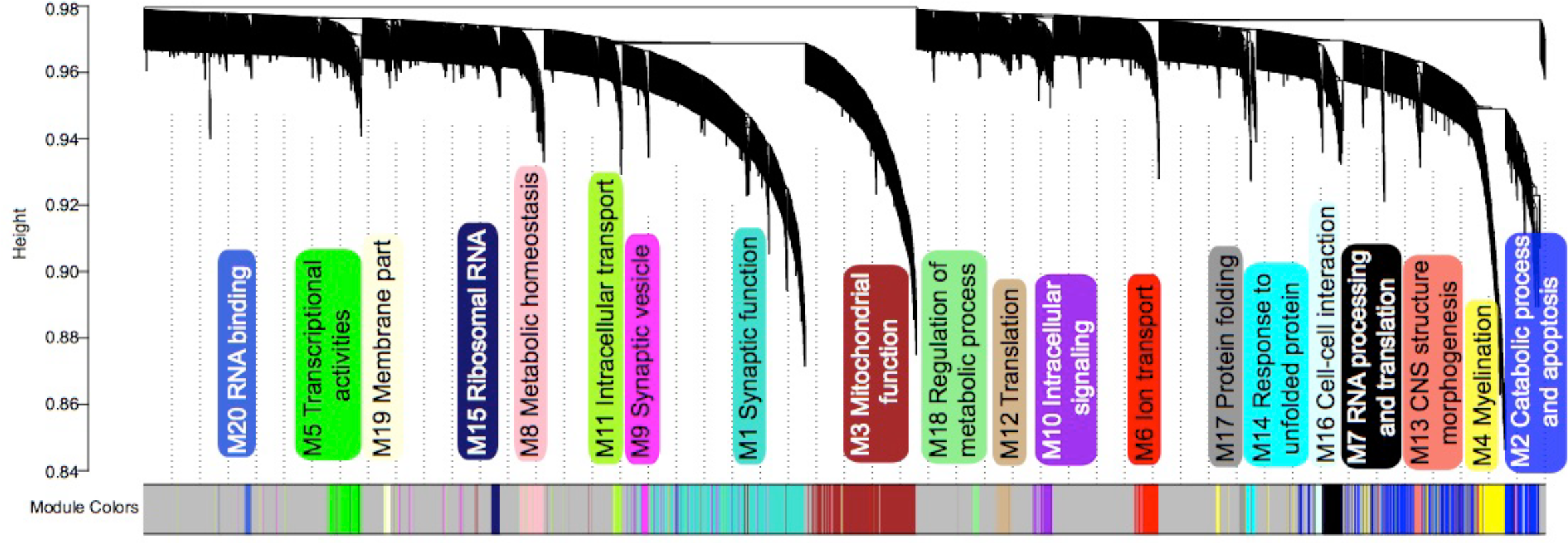
WGCNA cluster dendogram groups proteins into 20 distinct protein co-expressed modules defined by dendogram branch cutting in the Banner cohort.

### Cognitive trajectory-associated proteins and their relation to protein networks

The 579 proteins associated with cognitive trajectory from the meta-analysis were tested for enrichment in each network of co-expressed proteins in the Banner and BLSA cohorts, respectively. We considered the 350 higher-abundance proteins and the 229 lower-abundance proteins in cognitive stability separately. The higher-abundance proteins in cognitive stability were enriched in modules involved in synaptic function (Banner M1, BLSA M1 and M4) and mitochondrial function (Banner M3 and BLSA M3) in both cohorts at adjusted p <0.05 (Figure 5). On the other hand, the lower-abundance proteins in cognitive stability were enriched in the modules involved in apoptosis (Banner M2, BLSA M13), myelination (Banner M4, BLSA M2), and inflammatory response (BLSA M6 and indirectly Banner M2 by virtue of overlap) in both cohorts at adjusted p <0.05 (Figure 5).

**Figure 5:**
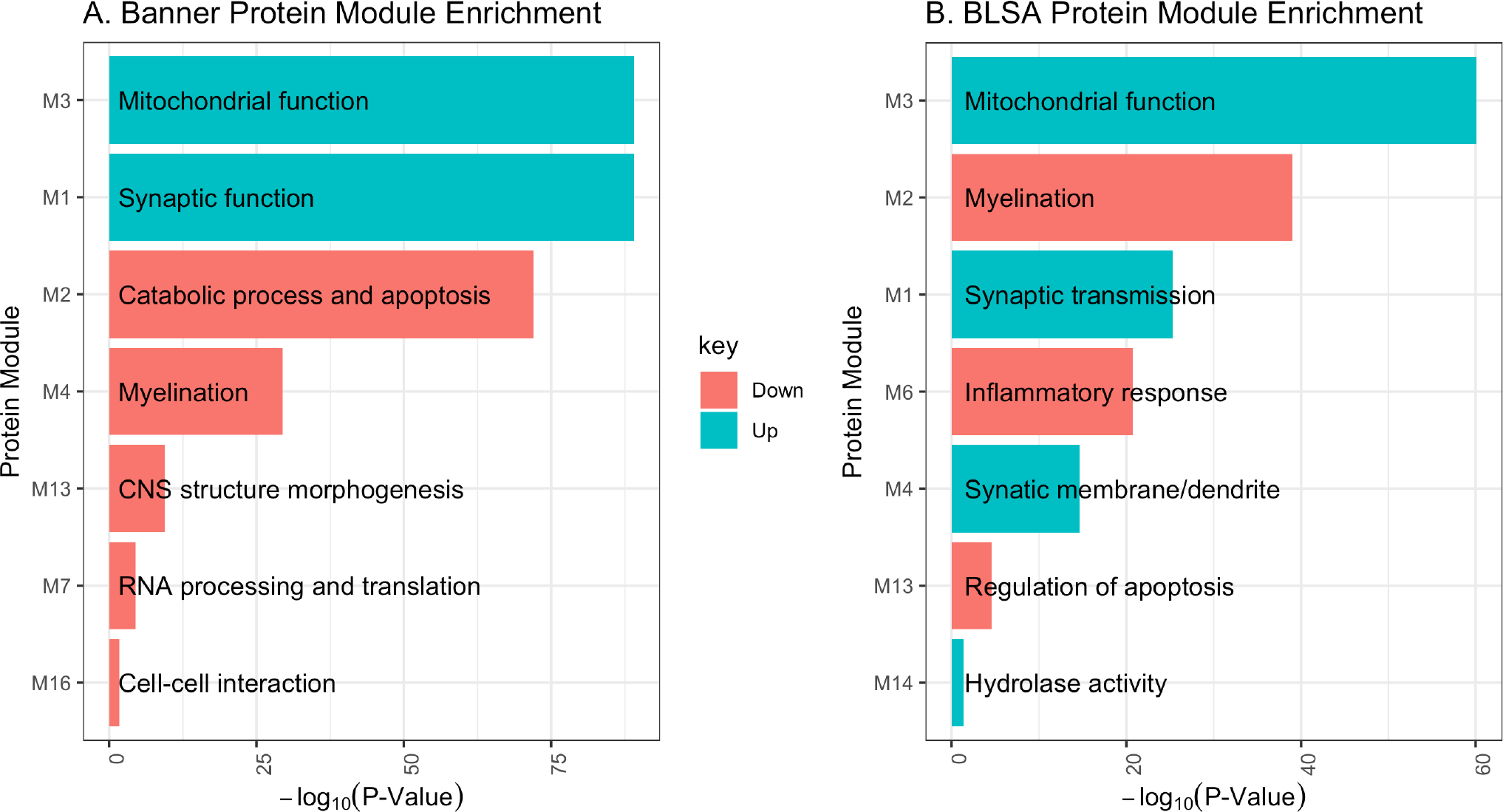
Enrichment of the 579 cognitive trajectory-associated proteins in modules of co-expressed proteins in Banner and BLSA cohorts. Enrichment analysis was performed separately for lower-abundance proteins in cognitive stability (229 proteins, colored in salmon) and higher-abundance proteins in cognitive stability (350 proteins, colored in turquoise), respectively. The P-values are FDR adjusted p-values. **a)** Banner protein module enrichment; **b)** BLSA protein module enrichment.

Among the 232 proteins associated with cognitive trajectory independently of plaques and tangles, we found that the higher-abundance proteins in cognitive stability were enriched for modules of synaptic function (Banner M1, BLSA M1 and BLSA M4), and mitochondrial function (Banner M3 and BLSA M3) in both cohorts at adjusted p <0.05 (Figure 6). The lower-abundance proteins in cognitive stability were enriched for the modules involving apoptosis (Banner M2, BLSA M13), myelination (Banner M4, BLSA M2), and inflammatory response (BLSA M6 and Banner M2 by virtue of overlap) in both cohorts at adjusted p <0.05 (Figure 6).

**Figure 6:**
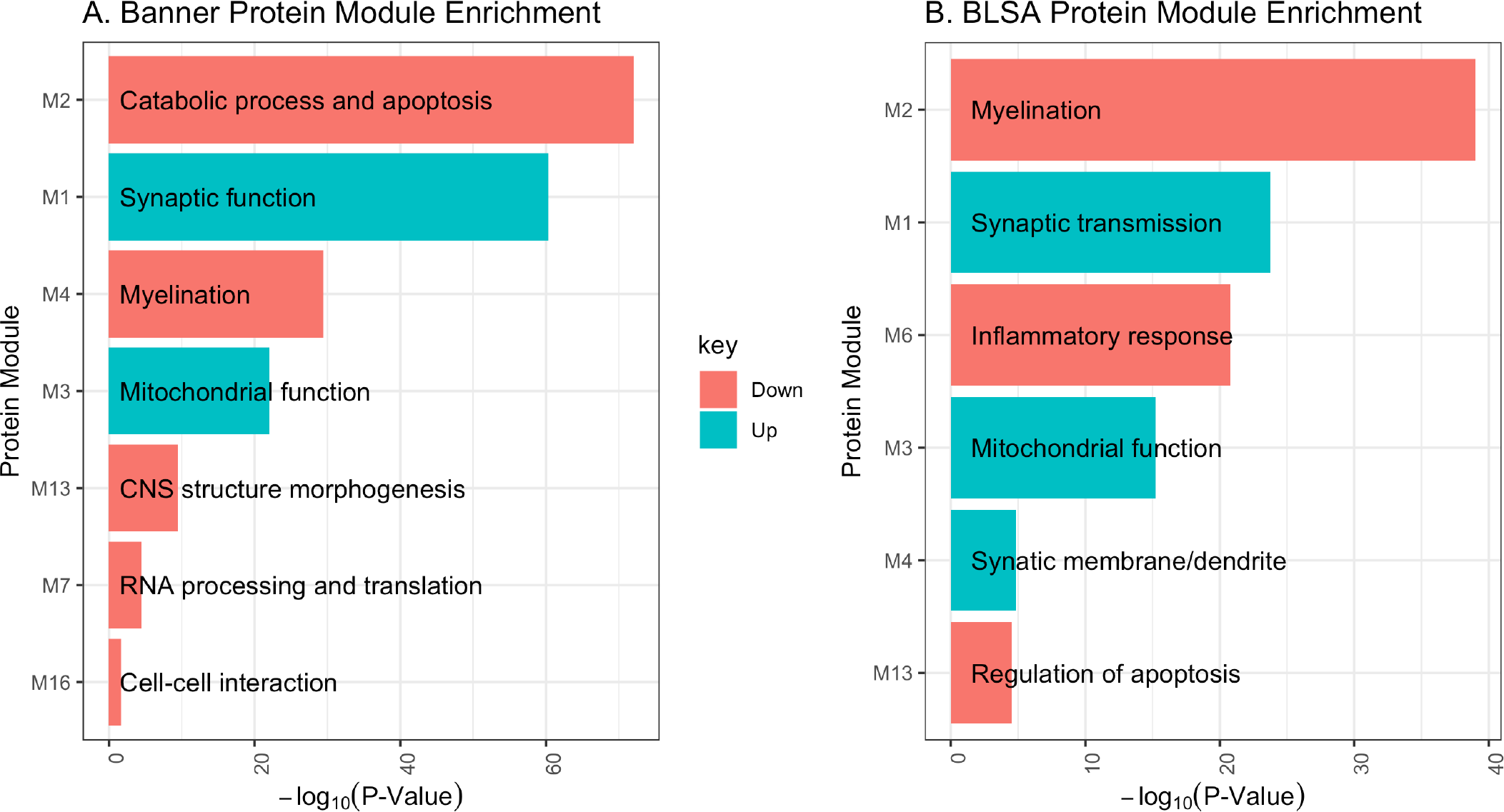
Enrichment of the 232 cognitive trajectory-associated proteins in modules of co-expressed proteins. Of note, these 232 proteins were associated with cognitive trajectory independently of plaques and tangles. Enrichment analysis was performed separately for lower-abundance proteins in cognitive stability (81 proteins, colored in salmon) and higher-abundance proteins in cognitive stability (151 proteins, colored in turquoise), respectively. The P-values are FDR adjusted p-values. **a)** Banner protein module enrichment; **b)** BLSA protein module enrichment.

We note that through both approaches - proteome-wide association study of cognitive trajectory and protein co-expression networks - we found consistent results that the over-expressed proteins in cognitive stability were enriched for mitochondrial activities and synaptic function independently of amyloid plaques and neurofibrillary tangles.

### Modules of co-expressed proteins and cognitive trajectory

We examined direct correlation between the modules of co-expressed proteins and cognitive trajectory in each cohort. In the Banner network, 10 of its 20 modules were associated with cognitive trajectory at FDR <0.05 (Table 4). The top six of these 10 modules were the same modules found to be enriched among the proteins associated with cognitive trajectory (Table 4, Figure 5A). In BLSA, four modules were associated with cognitive trajectory at p<0.05 but only one remained significantly associated with cognitive trajectory after adjusting for multiple testing (M3/mitochondrial function, p=0.0025, adjusted p = 0.039), likely due to small sample size and thus lower power (Table 4).

**Table 4:** Correlations between modules of co-expressed proteins and cognitive trajectory. Positive correlation indicates that higher abundance of the proteins in the module was associated with cognitive stability. On the other hand, negative correlation suggests that lower abundance of the proteins in the module was associated with cognitive stability. The table is sorted by correlation coefficient r and then by p-value. Modules not listed are not significantly associated with cognitive trajectory.

| Module | Biological process | $r$ | $p$ | Adjusted $p$ |
| --- | --- | --- | --- | --- |
| **BANNER (N=104)** |  |  |  |  |
| M1 | Synaptic function | 0.52 | 1.52E-08 | 1.52E-07 |
| M3 | Mitochondrial function | 0.39 | 3.73E-05 | 0.00019 |
| M2 | Catabolic process and apoptosis | -0.59 | 4.15E-11 | 8.30E-10 |
| M13 | CNS structure morphogenesis | -0.41 | 1.45E-05 | 9.66E-05 |
| M4 | Myelination | -0.34 | 0.000329 | 0.00132 |
| M7 | RNA processing and translation | -0.32 | 0.000847 | 0.00282 |
| M14 | Response to unfolded protein | -0.26 | 0.00758 | 0.0217 |
| M16 | Cell-cell interaction | -0.24 | 0.0121 | 0.0270 |
| M6 | Ion transport | -0.24 | 0.0144 | 0.0288 |
| M11 | Intracellular transport | 0.25 | 0.0097 | 0.0243 |
| **BLSA (N=35)** |  |  |  |  |
| M3 | Mitochondrial function | 0.50 | 0.0025 | 0.039 |
| M4 | Synaptic membrane/dendrite | 0.40 | 0.0180 | 0.085 |
| M6 | Inflammatory response | -0.39 | 0.0189 | 0.085 |
| M8 | Ribonucleoprotein complex | -0.39 | 0.0214 | 0.085 |

$r$ : correlation coefficient

After adjusting for amyloid plaques and neurofibrillary tangles, only two of the Banner modules M1/Synaptic function (β = 0.039, p = 6.74E−06, adjust p = 0.00013) and M2/Catabolic processes and apoptosis (β = −0.032, p = 3.02E−05, adjust p = 0.00030) remained significantly associated with cognitive trajectory after multiple testing adjustment at FDR <0.05. For BLSA, only one module, M3/mitochondrial function (β = −0.097, p = 0.0059, adjust p = 0.095), was associated with cognitive trajectory at adjusted p-value <0.1 after adjusting for plaques and tangles.

### Cognitive trajectory-associated proteins and hub proteins

In protein co-expression networks, hub proteins are likely important proteins because they are highly correlated with other proteins in the module. We sought to determine whether proteins associated with cognitive trajectory were also hub proteins in the Banner and BLSA co-expression networks. Hub proteins were defined as proteins with intramodular kME in the top 90^th^ percentile. We focused on the seven Banner and seven BLSA modules that the 579 cognitive trajectory-associated proteins were enriched in (Figure 5). In Banner, these seven modules had 123 hub proteins, of which 102 (83%) were associated with cognitive trajectory (Supplementary Table 15a). In BLSA, among the 131 hub proteins in the seven modules, 99 (76%) were associated with cognitive trajectory (Supplementary Table 15b). We performed the same analysis, except that the input proteins were those that associate with cognitive trajectory independently of pathologies. We found that among the 123 hub proteins in Banner and 127 hub proteins in BLSA, 67 and 62 respectively were also associated with cognitive trajectory independently of plaques and tangles (Supplementary Tables 16a,b). In common between these 67 (Banner) and 62 (BLSA) hub and cognitive trajectory-associated proteins were 38 proteins (Table 5, Supplementary table 17). Notably, these 38 proteins were associated with cognitive trajectory independently of amyloid plaques and neurofibrillary tangles in both Banner and BLSA, and were also hub proteins in both Banner and BLSA and thus are tractable targets for further mechanistic studies of cognitive stability.

**Table 5:**
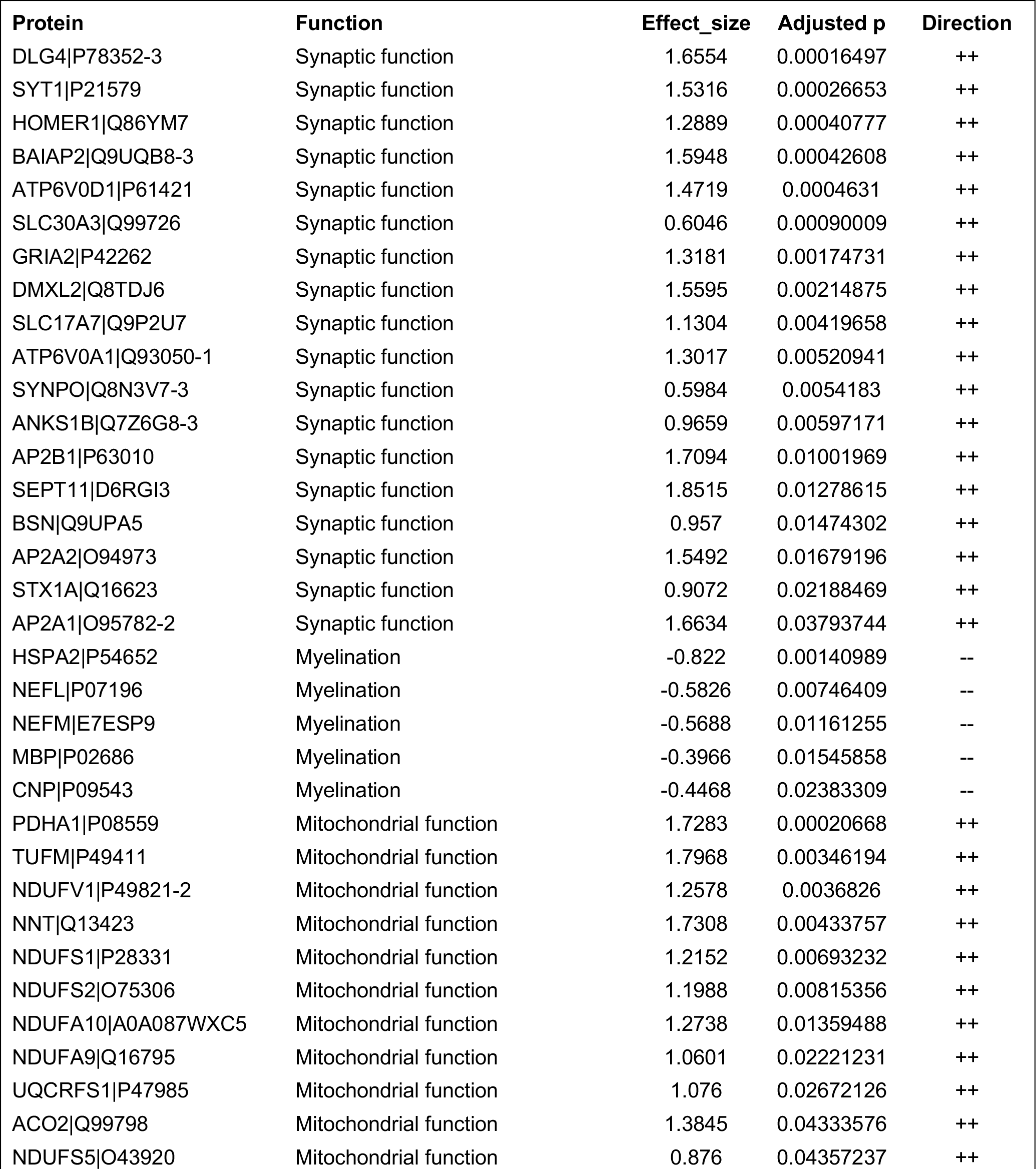
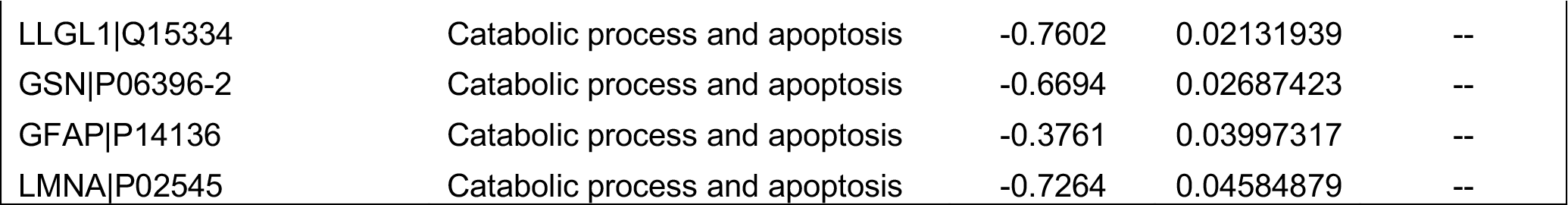
List of 38 hub proteins that were also associated with cognitive trajectory after adjusting for amyloid plaques and neurofibrillary tangles in both the discovery and replication cohorts. The proteins were sorted by the GO terms of the corresponding co-expression protein modules and then by adjusted p-value from the meta-analysis of proteome-wide association study of cognitive trajectory. Direction indicates high-abundance (++) or low-abundance (−−) in cognitive stability, respectively, in both the discovery and replication cohorts. Detailed information about these proteins are presented in Supplementary Table 17.

Remarkably, the top protein of this 38-protein list is DLG4, also known as PSD95, which is one of the most abundant scaffold proteins at the excitatory brain synapses and known to affect brain development, neuronal plasticity, memory and learning, and overall cognitive function^41–43^. We asked whether proteins known to interact with PSD95, i.e. PSD95 interactome, were also associated with cognitive trajectory using the published PSD95 interactome from Fernandez et al^41^. Among the 118 proteins known to interact with PSD95, 109 were detected in the Banner cohort. The first principal component of these 109 proteins, which captures 99.3% of the variance of the PSD95 interactome, was significantly associated with cognitive trajectory (β = 0.016; p = 0.0001; N=104; Supplementary Figure 6). This implies that higher expression of the PSD95 interacting partners was associated with cognitive stability (Supplementary Figure 6).

## DISCUSSION

We performed an unbiased proteome-wide association study of cognitive trajectory in a discovery and replication cohort, followed by a meta-analysis, to identify new biological processes underlying cognitive stability. Particularly, we used both proteome-wide differential analysis and protein co-expression network analysis to gain new insights into changes in individual proteins as well as networks of proteins in cognitive trajectory. Our most notable finding is that proteins involving mitochondrial activities or synaptic functions had increased abundance among individuals with cognitive stability regardless of the burden of amyloid plaques or neurofibrillary tangles. In other words, cognitive stability is associated with increased mitochondrial and synaptic activities independently of the burden of plaques or tangles in neurons predominantly. These findings were derived from two approaches in two different cohorts: i) proteome-wide association study of cognitive trajectory followed by GO enrichment analysis of the associated proteins, and ii) protein co-expression network enrichment analysis of the cognitive trajectory-associated proteins. Most interestingly, mitochondrial activities are strikingly present in our analysis in a way not seen when considering clinical diagnosis of cognitive status or AD pathologic outcomes. For instance, BLSA module M3, enriched for mitochondrial function, was most significantly associated with cognitive trajectory both before and after adjusting for plaques and tangles. This module was not significantly associated with clinical diagnosis or AD pathologies after multiple testing correction in our previous publication^14^, suggesting that cognitive trajectory captures new biological processes that are independent of traditional AD pathologies.

Mitochondria in neurons are crucial for maintaining synaptic function, and most of the mitochondrial proteins are encoded in the nuclear genome^44^. In the brain, mitochondria regulate synaptic transmission by generating ATP to power the process and by modulating presynaptic calcium level, which in turn determines the release of neurotransmitters^44^. Synaptic mitochondria, therefore, are vital for the maintenance of synaptic function and transmission^44–46^. This is consistent with our finding that 157 proteins of the 579 cognitive trajectory-associated proteins are mitochondrial proteins and of these, 122 are located in the mitochondria either in the pre-or postsynaptic density. Mitochondrial dysfunction has been suggested to occur early in the progression to AD and continues into later stages of AD^46–49^. An example of early manifestation of mitochondrial dysfunction is a study of expired young-adult *APOE4* carriers versus expired young-adult non-*APOE4* carriers^47^. In that study, young-adult participants in both groups did not have amyloid or tangle deposition in the posterior cingulate cortex (PCC); yet, *APOE4* carriers, i.e., those at greater risk for developing AD, showed reduced mitochondrial activity in PCC neurons, as measured by cytochrome oxidase levels, suggesting that a change in mitochondrial function may be the earliest manifestation of AD risk^47^. An example of mitochondrial dysfunction continuing into later stages of AD is a transcriptomic study of post-mortem brain of AD cases and controls in the PCC and other brain regions^48^. This study found that the nuclear genes encoding the mitochondrial electron transport chain subunits had decreased expression in AD cases compared to controls in PCC and other brain regions^48^. Furthermore, a transgenic mouse model showed that amyloid pathology, even at a low level, can induce deficits in the functioning of synaptic mitochondria leading to synaptic degeneration^45^. Synaptic degeneration has been suggested to be one of the earliest cellular event in AD pathogenesis, and synaptic loss to be the best correlate of cognitive impairment in AD (see review by Reddy et al^46^). These findings are consistent with ours, in which proteins involved in mitochondrial activities had lower abundance in persons with faster cognitive decline but higher abundance in persons with cognitive stability. To the best of our knowledge, ours is the first human unbiased proteome-wide study that shows greater abundance of mitochondrial proteins being linked to cognitive stability. Taken together, our findings and others highlight that mitochondrial activities would be a fruitful research target for early prevention of cognitive decline and enhancement of cognitive stability.

Synaptic function in cognitive decline has been given more attention than mitochondrial function. For instance, a study of candidate synaptic proteins in cognitive decline in AD found that reduced abundance of the synaptic protein SNAP25 was associated with faster cognitive decline^50^. Consistently, in both Banner and BLSA, we found that lower abundance of SNAP25 was associated with faster cognitive decline. Furthermore, Bereczki and colleagues performed a comprehensive study of 851 synaptic proteins in neurodegenerative diseases and cognitive decline^51^. They found that reduced levels of synaptic proteins SNAP47, SYBU, LRFN2, SV2C, and GRIA3 were associated with cognitive impairment and faster cognitive decline^51^. Among these proteins, only GRIA3 was detected in our datasets and had lower abundance in faster cognitive decline in both Banner and BLSA, consistent with Bereczki and colleagues’ finding.

The next notable finding is the 38 proteins that were associated with cognitive trajectory independently of typical AD neuropathologies and were also hub proteins in both Banner and BLSA protein networks. These proteins are highly promising targets for future mechanistic studies of cognitive trajectory. Among these 38 proteins, those that are higher abundance in cognitive stability are involved in either mitochondrial activities or synaptic function, and those with lower abundance in cognitive stability are involved in myelination or apoptosis. Among the proteins involved in synaptic function is DLG4, also known as PSD-95 (postsynaptic density-95 protein). PSD-95 is a major scaffold protein of the dendritic spines and its expression level has been shown to be altered in aging, AD, and several psychiatric disorders^52^. PSD-95 is important for neuronal plasticity and memory consolidation via its regulation of synaptic glutamate receptors, signaling proteins, adhesion molecules, and cytoskeletal proteins^52 53^. Here we show that the aggregate of PSD-95 interactome abundance is also associated with cognitive trajectory. Bustos and colleagues showed that increased protein level of PSD95 in the hippocampus rescued memory deficits of aged mice and dementia mice^53^. Their finding is consistent with ours in which over-expression of PSD95 protein was associated with cognitive stability. The second protein on the list, SYT1, was found to have reduced abundance in AD^54^. Overexpression of Syt1 in mouse hippocampi was found to promote “protective” presenilin-1 conformation^54^, consistent with our finding of its protective effect. Another protein on this list, HOMER1 (i.e., HOMER1a) is an important protein for synaptic plasticity and memory consolidation because it drives homeostatic scaling-down of excitatory synapses during sleep to remodel synapses and consolidate contextual memory^55^. Consistently, we found that protein level of HOMER1 was increased in cognitive stability. SLC30A3, aka ZNT3, is a synapse-specific vesicular zinc transporter that is important for zinc homeostasis and required for hippocampus-dependent memory^56^. Consistently, we found that SLC30A3 protein level was increased in cognitive stability. SLC17A7, aka VGLUT1, is a sodium-dependent phosphate transporter that is specifically expressed in the neuron-rich regions of the brain and transports glutamate. Decreased abundance of this protein can lead to susceptibility to neuroinflammation and disruption of synaptic plasticity^57^. Along the same theme, SYNPO has been found to be important for synaptic plasticity^58 59^. A notable theme among the proteins participating in synaptic function on this list of 38 proteins is that they are important for neuronal plasticity and memory consolidation and their over-expression is associated with cognitive stability.

Some proteins, including NEFM and MBP, participate in myelination on this list of 38 proteins. Myelination refers to the formation of a membranous sheath surrounding axons to increase signal transmitting speed between neurons and is not limited to early development but occurs throughout adulthood^60^. NEFM are neurofilaments that maintain neuronal caliber and participate in intracellular transport to axons and dendrites. NEFM was shown to accumulate in the hippocampus in the presence of amyloid pathology^61^. To retard human aging and age-related diseases, inhibition of insulin-like growth factor 1 (IGF1) has been proposed, and several studies have shown that neuron-specific deletion of IGF1R confers neuroprotection and improves behavior in AD transgenic mice^61–63^. From a genome-wide screen, a prominent observation after genetic ablation of IGF1 receptor in AD mice is the reduction in expression level of NEFM, down to control level, suggesting that lower abundance of NEFM is protective^61^. These findings are consistent with ours, in which lower abundance of NEFM was associated with cognitive resilience. Myelin basic protein, MBP, can influence beta-amyloid accumulation in the brain. Indeed, deletion of MBP gene led to a significant reduction in cerebral beta-amyloid levels in Tg-3xFAD mice^64^, consistent with our finding of decreased abundance of MBP being associated with cognitive stability.

Besides the 38 proteins above, we also presented i) 579 proteins associated with cognitive trajectory from our meta-analysis, ii) 232 proteins associated with cognitive trajectory independently of amyloid plaques and neurofibrillary tangles, iii) 102 proteins and 99 proteins associated with cognitive trajectory and were also hub proteins in Banner networks and BLSA networks, respectively. Furthermore, we note that VGF is strongly associated with cognitive trajectory with or without adjusting for AD neuropathology. VGF is also a hub protein for Banner M1 module, which involves synaptic function. Our findings suggest that VGF likely act through mechanisms independent of amyloid plaques and neurofibrillary tangles in contributing to cognitive decline. VGF is a neuropeptide that promotes hippocampal neurogenesis, dendritic maturation, and synaptic activity^65^. VGF is induced by BDNF and serotonin and is upregulated by antidepressants, exercise, and is reduced in animal models of depression^66^. VGF is another promising candidate for mechanistic study of cognitive stability, consistent with findings from this multiscale causal network modeling of AD^67^.

From the protein co-expression network analysis, we found 10 Banner protein modules and 1 BLSA protein module associated with cognitive trajectory. Fewer networks remained associated with cognitive trajectory after adjusting for plaques and tangles, suggesting that some networks act through pathology while others act independently of pathology to contribute to variation in cognitive trajectory. We also note that the published BLSA protein co-expression modules used here were constructed using the protein expression levels from both the dorsolateral prefrontal cortex (dPFC) and precuneus so the module memberships might be slightly different than those from the dPFC alone. On the other hand, proteins and protein co-expression networks that are consistently changed within the two regions likely reflect biologically meaningful differences.

We found cognitive trajectory-associated proteins to be enriched in astrocytes, which perform important roles for neural functioning. Astrocytes provide neurotrophic support to promote neuronal survival, are important in the formation and maturation of synapses, help control neurotransmitter concentrations, and play a role in maintaining the blood-brain barrier^68^. We found that both higher- and lower-abundance proteins in cognitive stability were enriched in astrocytes. Our findings may reflect the two types of astrocytes as described in the literature - reactive astrocytes that lose the ability to promote neuronal survival, outgrowth, and synaptogenesis, and induce death of neurons and oligodendrocytes, and homeostatic astrocytes that promote neural functioning^69^.

Our findings should be interpreted in light of the strengths and limitations of this study. First, this is an association study and thus it precludes causal inference. However, we nominated the tractable targets for further mechanistic studies of cognitive trajectory. Second, our label-free proteomic method yielded approximately 3900 proteins in the proteome, which is not as comprehensive of a proteomic profile as would be ideal, and larger proteomic analysis will undoubtedly add refinement to the proteins and protein networks involved in cognitive trajectory. Third, the available cognitive data was from the MMSE, which is widely used a screening tool that provides a coarse gauge of cognitive performance. Nevertheless, decline in MMSE generally parallels that of more detailed testing^70^. Fourth, our sample sizes are relatively small (n=104 and n=39) and exclusively Caucasians, which may limit the generalizability of our findings and power. Despite this, ours is the largest brain proteomic study of cognitive trajectory to our knowledge. These subjects were followed for a total of 887 person-years over 676 visits. Additional strengths of the study include that this is the first large scale proteome-wide study of cognitive trajectory to our knowledge. We also employed a discovery and replication cohort design followed by meta-analysis to reduce false positive findings. Furthermore, our focus on individual cognitive trajectory is more germane to early intervention and precision medicine. Finally, our findings establish a new framework for understanding mechanisms underlying cognitive trajectory and nominate a list of highly promising targets for future mechanistic studies of cognitive stability.

## ACKNOWLEDGMENTS

Support was provided by R01 AG056533, R01 AG053960, the Accelerating Medicine Partnership for AD (U01AG046161; U01 AG061357), the Emory Alzheimer’s Disease Research Center (P50 AG025688), and the NINDS Emory Neuroscience Core (P30 NS055077). This work was supported in part by funding from the intramural program of the National Institute on Aging (NIA). A.P.W. is also supported by U01 MH115484 and I01 BX003853. T.S.W. is also supported by IK2 BX001820 and P50 AG025688. N.T.S. is also supported by an Alzheimer’s Association, Alzheimer’s Research UK, The Michael J. Fox Foundation for Parkinson’s Research, the Weston Brain Institute Biomarkers Across Neurodegenerative Diseases Grant (11060), R01 AG061800, and R01 AG057911.

We are grateful to the Sun Health Research Institute Brain and Body Donation Program of Sun City, Arizona for the provision of human brain tissue. The Brain and Body Donation Program is supported by the National Institute of Neurological Disorders and Stroke (U24 NS072026 National Brain and Tissue Resource for Parkinson’s Disease and Related Disorders), the National Institute on Aging (P30 AG19610 Arizona Alzheimer’s Disease Core Center), the Arizona Department of Health Services (contract 211002, Arizona Alzheimer’s Research Center), the Arizona Biomedical Research Commission (contracts 4001, 0011, 05-901 and 1001 to the Arizona Parkinson’s Disease Consortium) and the Michael J. Fox Foundation for Parkinson’s Research.”

We are grateful to participants in BLSA for their invaluable contribution.

## AUTHOR CONTRIBUTIONS

A.P.W. and T.S.W. conceived and designed the experiments. D.M.D., J.C.T., M.T., T.G.B., G.E.S., E.M.R., R.J.C., J.J.L, N.T.S., and A.I.L. generated the experimental data. A.P.W., E.B.D., M.S.B., B.A.L., and T.S.W. performed the analyses. A.P.W., E.B.D., M.S.B., B.A.L., J.J.L., N.T.S., A.I.L., and T.S.W. interpreted the results. N.T.S., A.I.L., A.P.W., and T.S.W. acquired the funding. A.P.W. wrote the first draft of the manuscript. All authors revised and approved the manuscript.

## SUPPLEMENTARY DOCUMENTS

### SUPPLEMENTARY FIGURES

**Supplementary Figure 1:**
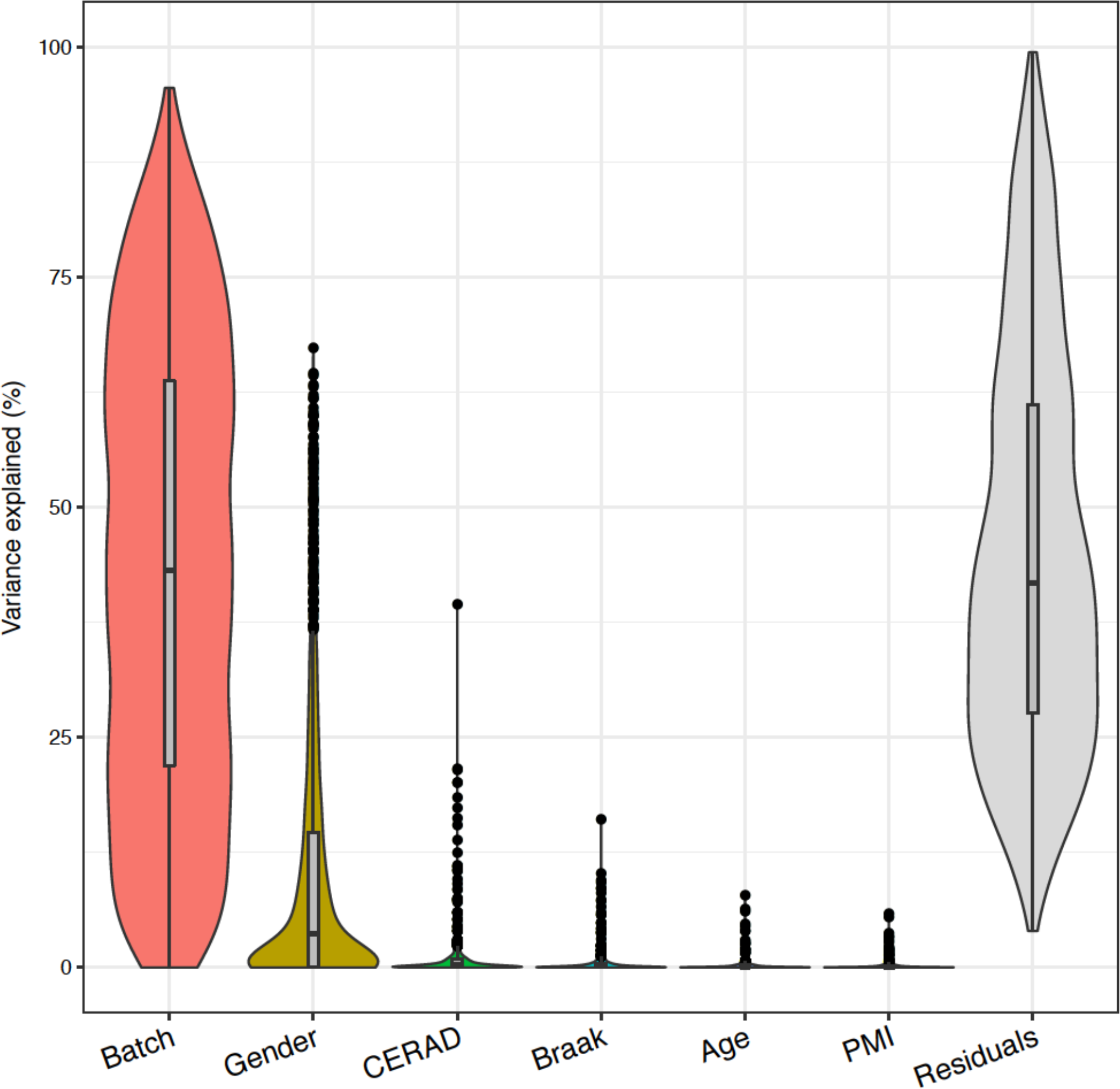
Variance partition of Banner proteomic profiles before regressing out effects of sex, age at death, post-mortem interval (PMI), and batch effects.

**Supplementary Figure 2:**
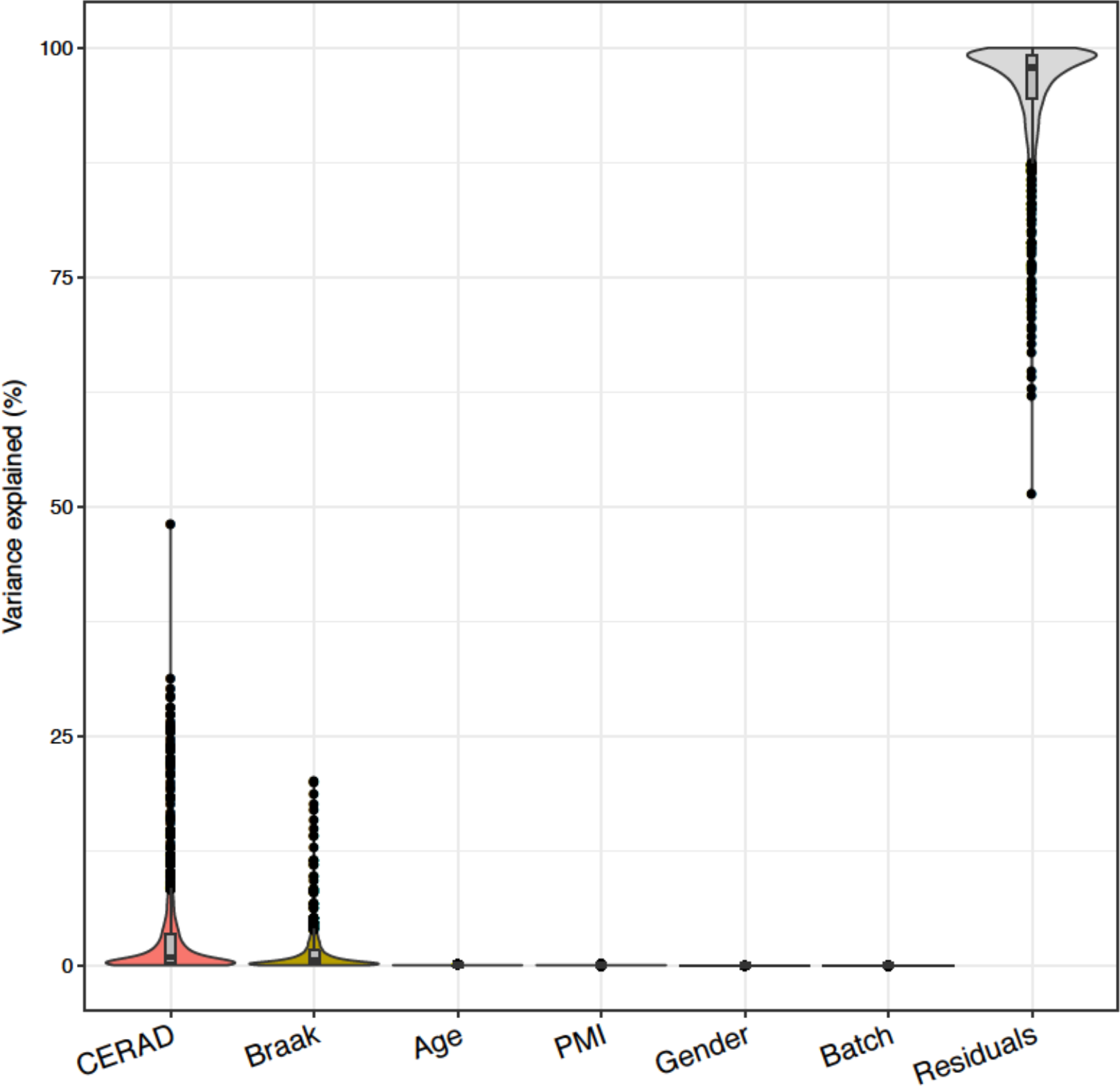
Variance partition of Banner proteomic profiles after regressing out effects of sex, age at death, post-mortem interval (PMI), and batch effects. This figure shows successful normalization of the proteomic profile.

**Supplementary Figure 3:**
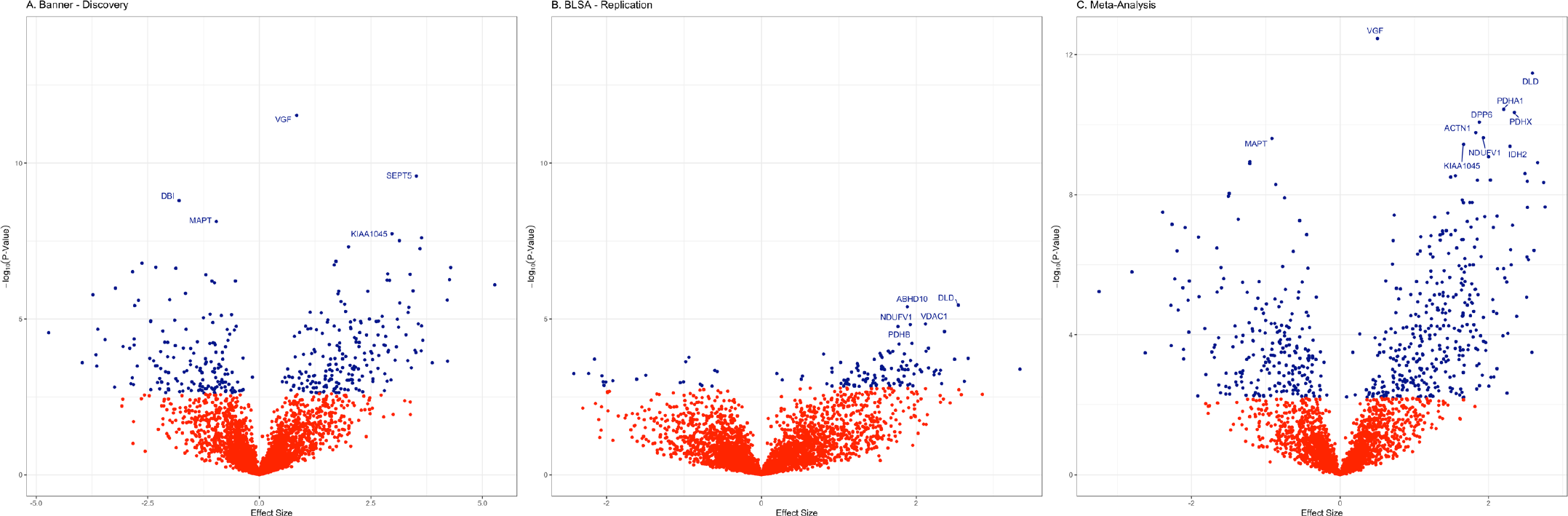
Volcano plot of proteome-wide association study of cognitive trajectory. The proteins significantly associated with cognitive trajectory were colored blue. **A.** Banner (Discovery) dataset with the top 5 proteins labeled. **B.** BLSA (Replication) dataset with the top 5 proteins labeled. **C.** Meta-analysis top 10 proteins labeled.

**Supplementary Figure 4:**
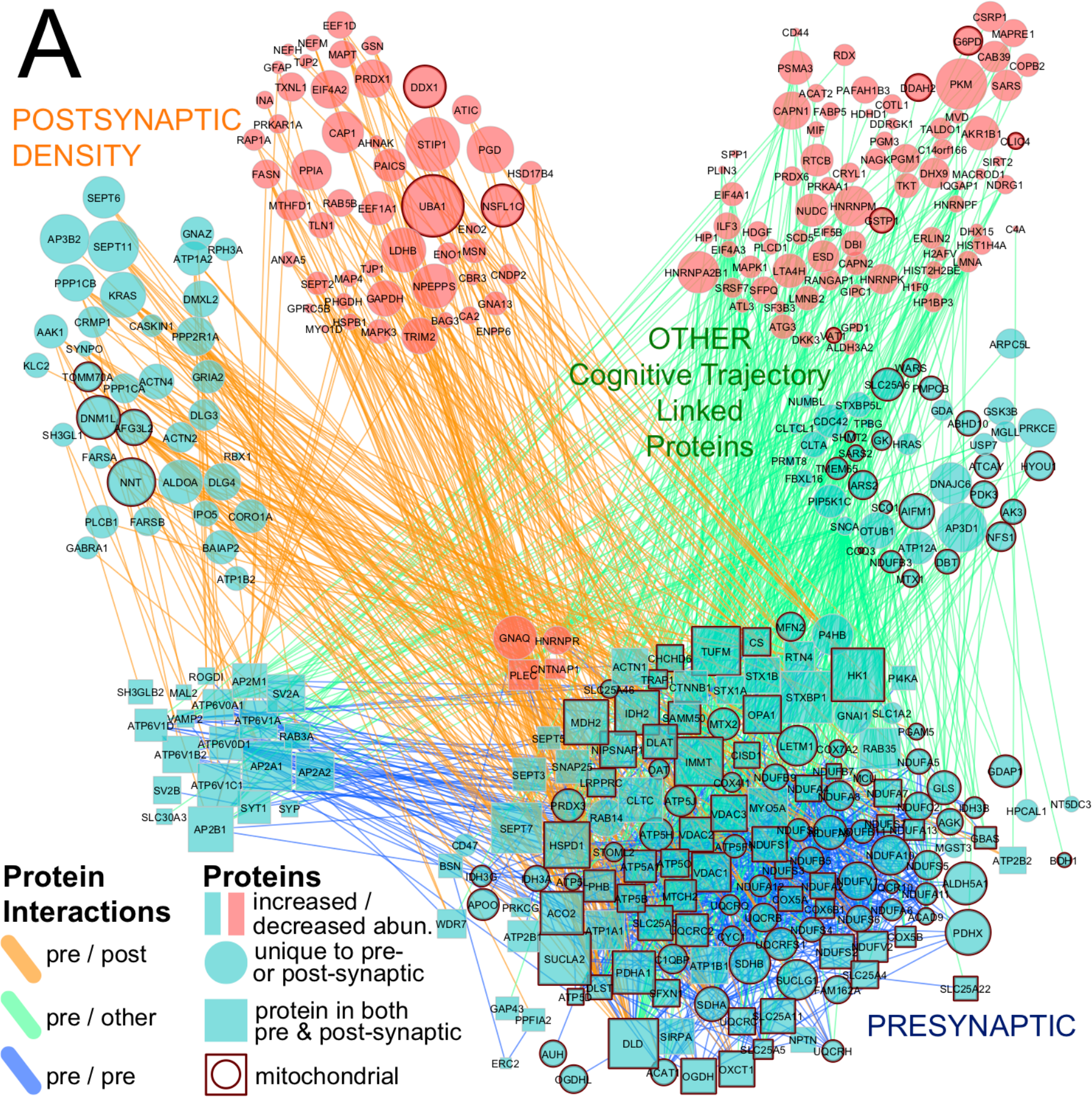
Cytoscape protein-protein interaction network based on BioGRID among the 579 proteins associated with cognitive trajectory. Among these 579 proteins, 569 were unique proteins. Among these 569 unique proteins, 159 were presynaptic, 150 postsynaptic, 20 present in both pre- and postsynaptic density, and 240 were located in other locations. Additionally, 112 presynaptic proteins, 10 postsynaptic proteins, and 35 other proteins were also designated as mitochondrial proteins.

**Supplementary Figure 5:**
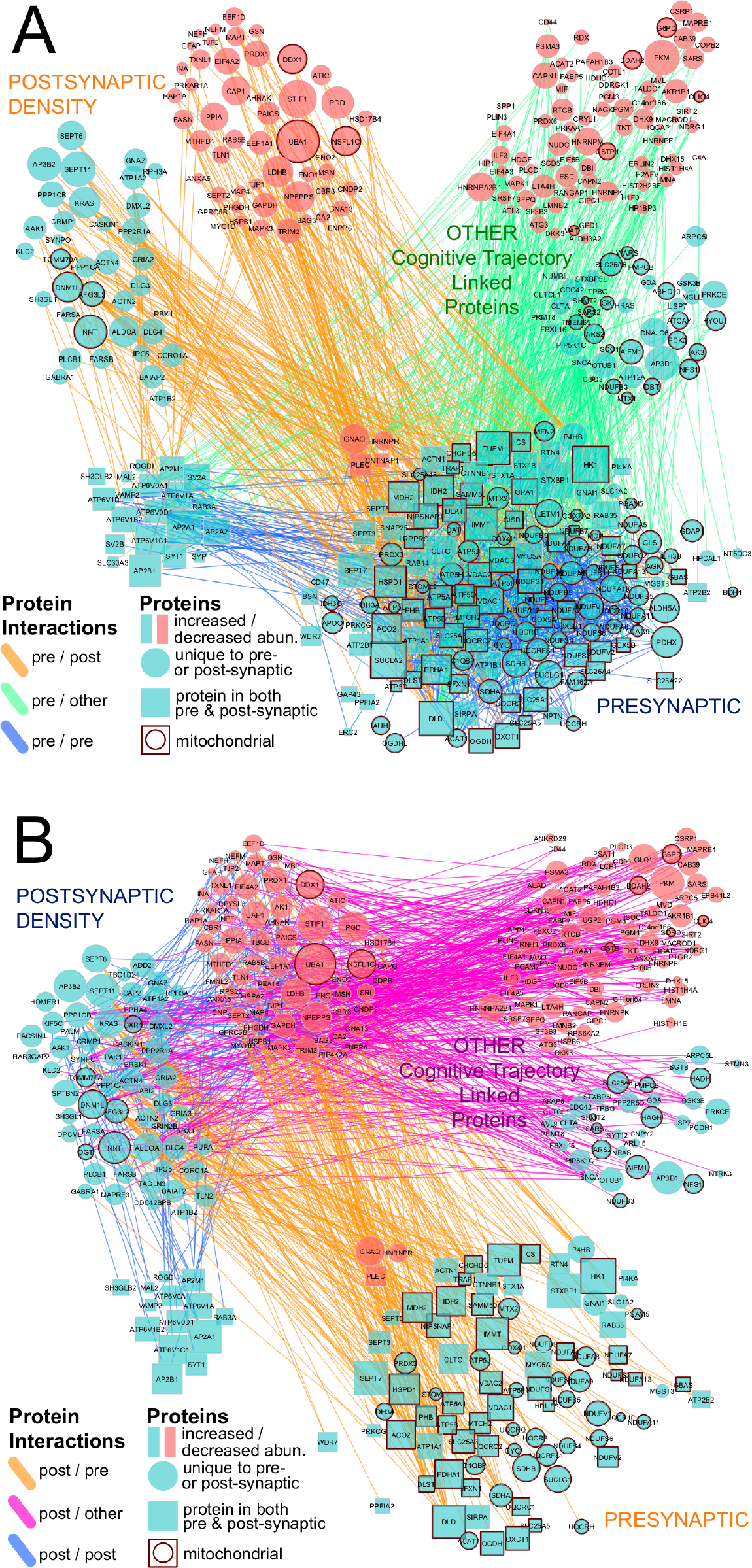
Cytoscape protein-protein interaction network based on BioGRID among the 579 proteins associated with cognitive trajectory. Among these 579 proteins, 569 were unique proteins. Among these 569 proteins, 159 were presynaptic, 150 postsynaptic, 20 present in both pre- and postsynaptic density, and 240 proteins were located in other locations. Additionally, 112 presynaptic proteins, 10 postsynaptic proteins, and 35 other proteins were also designated as mitochondrial proteins. **Panel A** focuses on the protein-protein interaction (PPI) emanating from presynaptic proteins. **Panel B** focuses on the PPI emanating from postsynaptic proteins.

**Supplementary Figure 6:**
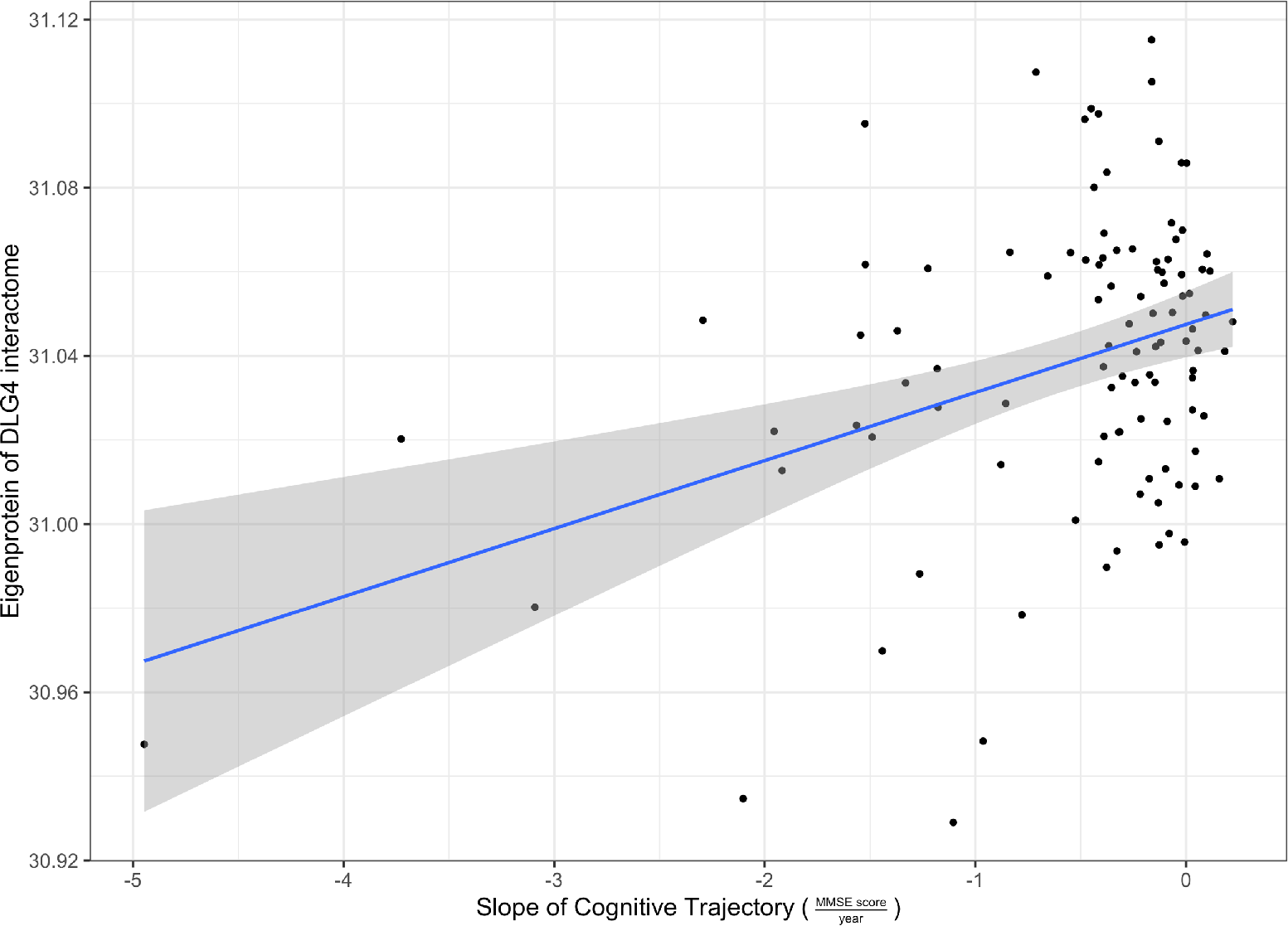
Interactome of DLG4/PSD95 and cognitive trajectory in Banner cohort. Higher expression of proteins of DLG4/PSD95 interactome was significantly associated with cognitive stability. The grey area denotes 95% confidence interval.

### SUPPLEMENTARY TABLES

**supplementary table 1:** Cognitive trajectory, number of MMSE tests done, and duration from last MMSE to death for each subject in Banner and BLSA cohorts.

**Supplementary Table 1a**: Proteome-wide association study of cognitive trajectory in the Banner discovery cohort adjusting for sex, age at enrollment, education, age at death, PMI, and batch effect (N=104).

**supplementary table 2**: Proteome-wide association study of cognitive trajectory in the BLSA replication cohort adjusting for sex, age at enrollment, education, age at death, PMI, and batch (N=39).

**supplementary table 3**: A meta-analysis of proteome-wide association studies of cognitive trajectory in the discovery and replication cohorts (N=143) revealed 579 proteins associated with cognitive trajectory in the same directions in both cohorts and at FDR p<0.05.

**supplementary table 3b**: Designation of presynaptic, postsynaptic, both (present in both presynaptic and postsynaptic density), or other location for the 579 proteins associated with cognitive trajectory. Additional designation of mitochondrial protein was also provided.

**supplementary table 3c:** Pairwise protein-protein interactions based on BioGRID for the 579 cognitive trajectory-associated proteins.

**supplementary table 4**: Gene ontology enrichment analysis of the 579 cognitive trajectory-associated proteins: 4a) GO Biological Process enrichment analysis for 229 low-abundance proteins in cognitive stability; 4b) GO Cellular Component enrichment analysis for 229 low-abundance proteins in cognitive stability; 4c) GO Molecular Function enrichment analysis for 229 low-abundance proteins in cognitive stability; 4d) GO Biological Process enrichment analysis for 350 high-abundance proteins in cognitive stability; 4e) GO Cellular Component enrichment analysis for 350 high-abundance proteins in cognitive stability; 4f) GO Molecular Function enrichment analysis for 350 high-abundance proteins in cognitive stability.

**supplementary table 5**: Brain cell type specific enrichment analysis for the 579 cognitive trajectory-associated proteins. Cell type specific signatures were from Sharma et. al., Zeisel et. al., and Darmanis et. al. Enrichment analysis was performed separately for the 229 low-abundance proteins and 350 high-abundance proteins in cognitive stability, respectively.

**Supplementary Table 6:** Proteome-wide association study of cognitive trajectory in the Banner discovery cohort adjusting for sex, age at enrollment, education, age at death, PMI, batch, and amyloid plaques and neurofibrillary tangles (N=104).

**Supplementary Table 7**: Proteome-wide association study of cognitive trajectory in the BLSA replication cohort adjusting for sex, age at enrollment, education, age at death, PMI, batch, amyloid plaques and neurofibrillary tangles (N=39).

**Supplementary Table 8**: A meta-analysis of proteome-wide association studies of cognitive trajectory in the discovery and replication cohorts, adjusting for plaques and tangles and other covariates (N=143), revealed 232 proteins associated with cognitive trajectory in the same directions in both cohorts at FDR p<0.05.

**Supplementary Table 9**: Gene ontology enrichment analysis of the 232 cognitive trajectory-associated proteins. Of note, these 232 proteins were associated with cognitive trajectory after adjusting for plaques, tangles, and other covariates: a) GO Biological Process enrichment analysis for 81 low-abundance proteins in cognitive stability; b) GO Cellular Component enrichment analysis for 81 low-abundance proteins in cognitive stability; c) GO Molecular Function enrichment analysis for 81 low-abundance proteins in cognitive stability; d) GO Biological Process enrichment analysis for 151 high-abundance proteins in cognitive stability; e) GO Cellular Component enrichment analysis for 151 high-abundance proteins in cognitive stability; f) GO Molecular Function enrichment analysis for 151 high-abundance proteins in cognitive stability.

**Supplementary Table 10**: Brain cell type specific enrichment analysis for the 232 cognitive trajectory-associated proteins after adjusting for plaques, tangles, and other covariates. We used cell type specific signatures from Sharma et. al., Zeisel et. al., and Darmanis et. al. Enrichment analysis was performed separately for the 81 low-abundance proteins and 151 high-abundance proteins in cognitive stability, respectively.

**Supplementary Table 11**: WGCNA analysis identified 20 modules of co-expressed proteins in Banner networks.

**Supplementary Table12**: Modules of co-expressed proteins and the biological processes they are enriched in in Banner networks.

**Supplementary Table 13**: GO enrichment analysis for modules of co-expressed proteins in Banner.

**Supplementary Table 14:** Overlap analysis of protein modules from Banner and BLSA networks. The table lists only pairs of modules with significant overlap after multiple testing correction.

**Supplementary Table 15: a)** List of proteins that were hub proteins for a particular module and were also associated with cognitive trajectory in Banner; **b)** List of proteins that were hub proteins for a particular module and were also associated with cognitive trajectory in BLSA. Hub proteins were defined as those with intramodular kME in the top 90^th^ percentile.

**Supplementary Table 16:** List of proteins that were hub proteins for a particular module and were also associated with cognitive trajectory after adjusting for plaques and tangles in **a)** Banner; **b)** BLSA. Hub proteins were defined as those with intramodular kME in the top 90^th^ percentile.

**Supplementary Table 17:** List of 38 proteins that were hub proteins in both Banner and BLSA networks and were also associated with cognitive trajectory after adjusting for plaques and tangles in both Banner and BLSA cohorts.

